# HIV-specific CD8^+^ T-cells in tonsils express exhaustive T_RM_-like signatures

**DOI:** 10.1101/2021.11.04.467061

**Authors:** Rabiah Fardoos, Sarah K. Nyquist, Osaretin E. Asowata, Samuel W. Kazer, Alveera Singh, Abigail Ngoepe, Janifer Giandhari, Ntombifuthi Mthabela, Dirhona Ramjit, Samita Singh, Farina Karim, Søren Buus, Frank Anderson, James Z. Porterfield, Andile L. Sibiya, Rishan Bipath, Kumeshan Moodley, Warren Kuhn, Bonnie Berger, Son Nguyen, Tulio de Oliveira, Thumbi Ndung’u, Philip Goulder, Alex K. Shalek, Alasdair Leslie, Henrik N. Kløverpris

**Author notes:** These authors contributed equally to this work (Rabiah Fardoos, Sarah K. Nyquist). Corresponding author Address correspondence to: Henrik N. Kløverpris, Africa Health Research Institute, K-RITH Tower Building, 719 Umbilo Road, Durban 4001, South Africa and University of Copenhagen, MAERSK tower building 07-13, Blegdamsvej 3B, 2200 Copenhagen N, Denmark.;45.29720910.

## Abstract

Lymphoid tissues are an important HIV reservoir site that persists in the face of antiretroviral therapy and natural immunity. Targeting these reservoirs by harnessing the antiviral activity of local tissue resident memory ( T_RM_) CD8^+^ T-cells is of great interest, but limited data exist on T_RM_s within lymph nodes of people living with HIV (PLWH). Here, we studied tonsil CD8^+^ T-cells obtained from PLWH and uninfected controls from South Africa. We show that these cells are preferentially located outside the germinal centers (GCs), the main reservoir site for HIV, and display a low cytolytic and transcriptionally T_RM_-like profile that is distinct from blood. In PLWH, CD8^+^ T_RM_-like cells are highly expanded and adopt a more cytolytic, activated and exhausted phenotype characterized by increased expression of CD69, PD-1 and perforin, but reduced CD127. This phenotype was enhanced in HIV-specific CD8^+^ T-cells from tonsils compared to matched blood. Single-cell profiling of these cells revealed a clear transcriptional signature of T-cell activation, clonal expansion and exhaustion *ex-vivo*. In contrast, this signature was absent from HIV-specific CD8^+^ T-cells in tonsils isolated from a natural HIV controller, who expressed lower levels of cell surface PD-1 and CXCR5, and reduced transcriptional evidence of T-cell activation, exhaustion and cytolytic activity. Thus, we show that HIV-specific T_RM_-like CD8^+^ T-cells in tonsils from non-HIV controllers are enriched for activation and exhaustion profiles compared to those in blood, suggesting that lymphoid HIV-specific CD8^+^ T_RM_ cells are potentially ideal candidates for immunotherapy to modulate their ability to targeting the HIV reservoirs.

## Introduction

No treatment for HIV infection is currently available to eradicate viral reservoirs, except anecdotal reports involving immune ablation followed by adoptive transfer of non-susceptible target cells [1, 2]. A key challenge is the presence of these viral reservoirs within human tissues, primarily within the germinal center (GC) of lymphoid tissue structures [3, 4], which persist in spite of highly effective antiretroviral treatment (ART) [5–9]. Even when ART is initiated during the acute phase of infection [10], these reservoirs can still fuel rapid rebound of plasma viremia during treatment interruption [11]. Therefore, HIV treatment will continue to require life-long ART, unless therapeutics can be developed to purge virus from these sanctuary sites. A potential to achieve this goal is to enhance pre-existing HIV-specific CD8^+^ T-cell responses within these sites and thereby facilitating the eradication of the viral reservoir. However, this requires a better understanding of the CD8^+^ T-cells present in HIV infected lymphoid tissues.

Tissue resident memory (T_RM_) CD8^+^ T-cells [12] are retained within tissues, including lymphoid tissues, and characterised by the expression of CD69, ↵E integrin (CD103) and absence of sphingosine-1-phosphate receptor (S1PR1) [13–16]. CD8^+^ T_RM_ cells are involved in the early response to pathogens [17, 18] and initiate rapid defense upon reinfection [19]. CD8^+^ T-cells mediate viral control and protect from disease progression in HIV and simian immunodeficiency virus (SIV)-infected rhesus macaque and play a significant role in restricting viral reservoirs within lymphoid tissue [20–22] also during ART [4, 23]. However, in the vast majority of cases, viral control eventually fails in the absence of ART, and CD8^+^ T-cell function becomes impaired [24–26].

One obstacle for optimal CD8^+^ T-cell function and their ability to clear viral reservoirs may be related to elevated expression of programmed cell death protein 1 (PD-1) and other co-inhibitory molecules [27–29]. In the blood, high levels of antigen exposure during chronic untreated HIV infection lead to increased CD8^+^ T-cell activation, terminal differentiation and ultimately dysfunction termed ‘exhaustion’ [30–35], which is directly linked to cognate epitope availability [33, 36, 37]. However, PD-1 expression is elevated on T-cell subsets within tissue sites such as lymphoid organs, and may not directly relate to immune exhaustion [38]. In addition, HIV-specific CD8^+^ T-cells identified in tissue typically lack the GC homing receptor CXCR5 expression [3, 39, 40], which may limit their ability ells to clear HIV reservoirs from the immune privileged B-cell follicles [41].

Most studies of T-cell function in people living with HIV (PLWH) are of cells derived from the blood, which do not capture the phenotype and function of CD8^+^ T_RM_ located in close proximity to the viral reservoirs within lymphoid tissues [3, 4, 8]. Recent studies show important differences between HIV-specific CD8^+^ T-cells located in blood and lymphoid tissues, and highlight the importance of better defining the functionality of the cells in closest proximity to viral reservoirs [38, 42–44]. In particular, little is known about the expression of inhibitory receptors, such as PD-1, on HIV-specific CD8^+^ T-cells and the functional significance of these markers within HIV-infected tissue sites [14, 38]. This information is important for strategies seeking to enhance antiviral activities in HIV-infected tissue.

Here, we studied CD8^+^ T-cells in blood and palatine tonsils, a lymphoid tissue organ in the oral-pharyngeal mucosa [45], from PLWH recruited from HIV endemic areas in South Africa. We combined *in situ* phenotyping, flow cytometry and applied paired single cell RNA (scRNA) and T-cell receptor (TCR) sequencing on bulk and antigen specific CD8^+^ T-cells (HIV and CMV) to determine the impact of HIV in this compartment. We show that HIV infection increases PD-1 expression along with canonical CD69 and CD103 T_RM_ markers on tonsil CD8^+^ T-cells and that HIV-specific CD8^+^ T_RM_-like cells are associated with a distinct transcriptional signature of activation and exhaustion. This appears to be driven by the presence of detectable HIV antigen and is highest in expanded HIV-specific TCR-clonotypes. By contrast, in one natural HIV controller studied, HIV-specific CD8^+^ T_RM_-like cells express reduced levels of PD-1, increased follicle homing chemokine receptor CXCR5 infection and upregulation of non-cytolytic T-cell activity, suggesting natural HIV controllers harbour distinct CD8^+^ T_RM_-like function within lymphoid tissue [43].

## Results

### CD8^+^ T-cells from human tonsils express a non-cytolytic T_RM_-like phenotype and are located outside the follicular GC

Recent studies show major phenotypic differences between CD8^+^ T-cells in circulation and those in lymphoid tissue [38, 44]. To further explore these differences and examine the impact of HIV infection, we collected tonsils and peripheral blood from study participants undergoing routine tonsillectomy in clinics located within high HIV endemic areas of KwaZulu-Natal, South Africa (Table 1). First, we analyzed matched blood and tonsils from HIV uninfected participants (Figure S1A) and, consistent with previous work, observed reduced expression of the key T-cell cytolytic markers perforin and granzyme B in tonsil CD8^+^ T-cells (Figure S1A, Figure 1A) [44]. Overall, tonsil CD8^+^ T-cells were phenotypically distinct from the blood compartment (Figure 1B, Figure S1B), with higher surface expression of CD69, CD103, CD27 and the follicle-homing chemokine receptor CXCR5 (Figure 1C), consistent with a T_RM_-like phenotype [38]. CD127 (IL-7Ra) and PD-1 expression also tended to be higher in the tonsils, but this difference was not statistically significant. Next, we performed single-cell RNA-sequencing (scRNAseq) on bulk CD8^+^ T-cells isolated from blood (n=3) and tonsils (n=2) using the Seq-Well S3 platform [46, 47] (Figure 1D). We identified a distinct transcriptional program in tonsil CD8^+^ T-cells, with 223 differentially expressed genes (DEGs) characteristic of tissue resident markers previously established for lymph nodes [48] (Figure 1E, Table S1). Canonical cytotoxic genes included *GNLY, GZMB, PRF1* and *GZMH* were enriched in blood, with the notable exception of *GZMK*, which was only expressed in tonsil CD8^+^ T-cells (Figure 1F). Transcription factors *TBX21, EOMES* and *RORA* also displayed compartment specific patterns, with *EOMES* enriched in tonsil CD8^+^ T-cells consistent with recent findings [49], and *TBX21* (T-bet) and *RORA* in blood (Figure 1F). *KLF2*, involved in Sphingosine-1-phosphate receptor (S1PR) expression, was associated with *S1PR1* and *S1PR5* expression, all of which were reduced in tonsil CD8^+^ T-cells, consistent with their role of sphingosine-1-phosphate-mediated egress from lymphoid tissue into circulation. In addition, tonsil CD8^+^ T-cells expressed lower levels of the integrin encoding genes *ITGB2* and *ITGA4* and a apparent complete absence of *ITGB1* (Figure 1F). Recent work suggest that CD8^+^ T-cell cytotoxic activity is closely associated with the expression of ITGB1 (CD29) [50] and thus the lack of this integrin in particular further supports the non-cytotoxic phenotype of LN CD8^+^ T-cells under normal conditions. Flourescence microscopy showed that tonsil CD8^+^ T-cells in HIV negative individuals are primarily found outside the follicular germinal centers (GCs), in contrast to CD4 T-cells, which are observed both in the cortex and within GCs (Figure 1G). Thus, profiling of CD8^+^ T-cells in blood and tonsil defines a distinct protein and transcription factor expression signature, consistent with reduced cytolytic activity [31], which is largely confined outside GCs.

**Table 1.**
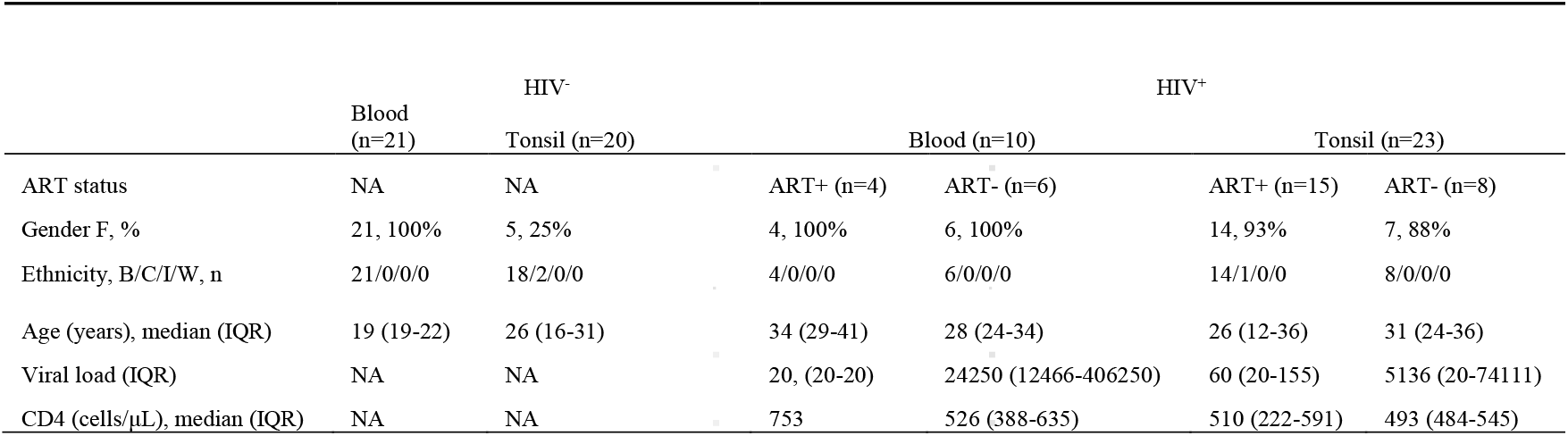
Clinical characteristics of study participants (n=64)

**Figure 1:**
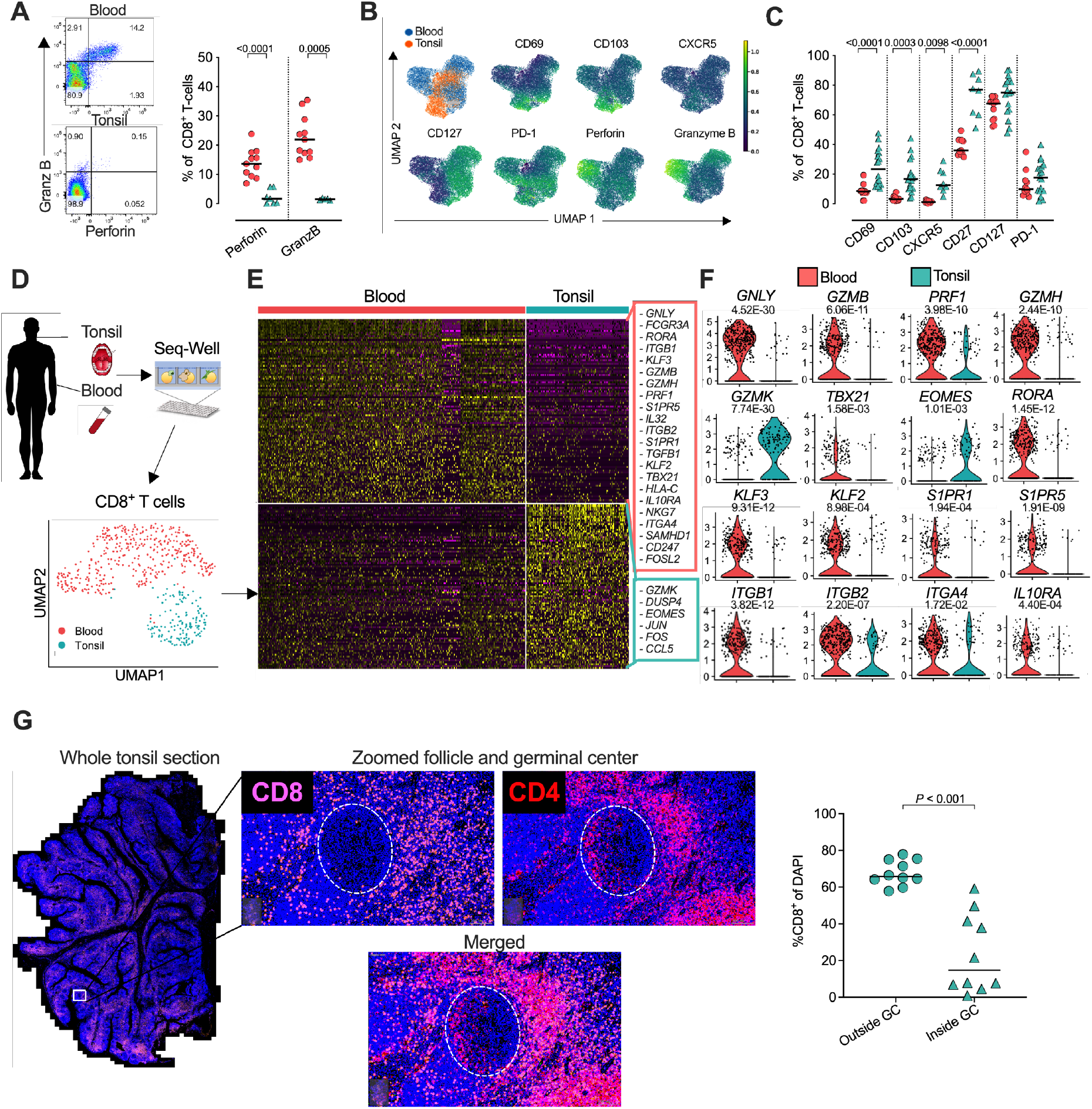
Blood and Tonsil CD8^+^ T-cell phenotype, single cell transcriptional profile and CD8^+^ T-cell in situ location within whole tonsil tissue sections. **A**. Blood and tonsil mononuclear cells pregated on CD8 T-cells by flow cytometry for perforin and granzyme B. **B**. Expression intensity of CD69, CD103, CXCR5, CD127, PD-1, Perforin and granzyme B in multidimensional tSNE space for CD8^+^ T-cells, gated from total CD45^+^ T-cells. Peripheral blood (blue), tonsil tissue (orange). **C**. Percentage of blood and tonsil CD8^+^ T-cells expressing indicated markers. **D**. Schematic of protocol for isolate of HIV^+^ CD8^+^ T cells from peripheral blood and tonsil tissue by scRNA-seq using Seq-Well v3 (top). UMAP of CD8^+^ T cells coloured by tissue source (bottom). **E**. Heatmap of z-scored gene expression of CD8^+^ T cells illustrating differentially expressed genes between blood and tonsil. Selective specifically expressed genes are marked alongside. **F**. Violin plots showing selected genes for cytolytic, transcription factors, tissue-resident memory and immune suppressive markers, expressed in CD8^+^ T cells. FDR-adjusted p<0.05; full results can be found in Supplementary Table S1. **G**. Fluorescent immunohistochemistry of whole tonsil section (left) from HIV uninfected donor shown by CD8, CD4 and merged panels (middle) with quantification using 10 unrelated areas identified outside follicles (outside GCs) and within follicular germinal centers ( inside GCs). P-values by Kruskal-Wallis multi comparisons.

### PD-1 expression and T_RM_-like phenotypes are elevated in tonsil CD8^+^ T-cells from PLWH

Next, we compared expression of phenotypic markers in CD8^+^ T-cells from tonsils and blood of PLWH (Figure S2A,B). The tissue resident markers CD69 and CD103 were upregulated in tonsils from PLWH, but not in matched blood from the same individuals, potentially indicating an enrichment of tonsil T_RM_ CD8^+^ T-cells after HIV infection (Figure 2A). CXCR5 was slightly elevated, though not statistically significant, potentially suggesting increased potential for accessing of CD8^+^ T-cells to the GC in PLWH through the CXCR5/CXCL13 axis. We found lower CD127, required for IL-7 homeostatic signalling, expression in both blood and tonsil. Conversely, PD-1 expression was highly upregulated in tonsil derived CD8^+^ T-cells of PLWH, suggesting antigen specific activation [51, 52] and potentially reduced functionality [33, 53]. GZMB expression was only elevated in the blood but not significantly in tonsils. However, perforin was significantly elevated in both compartments in PLWH suggesting a potential increase in cytolytic activity. Analysis of the co-expression of these markers in tonsil CD8^+^ T-cells indicates that HIV infection is associated with an increase in subsets expressing CD69^+^/CD103^+^/^−^ and PD-1^+^, and a reduced expression of CD127^+^ (Figure 2B).

**Figure 2:**
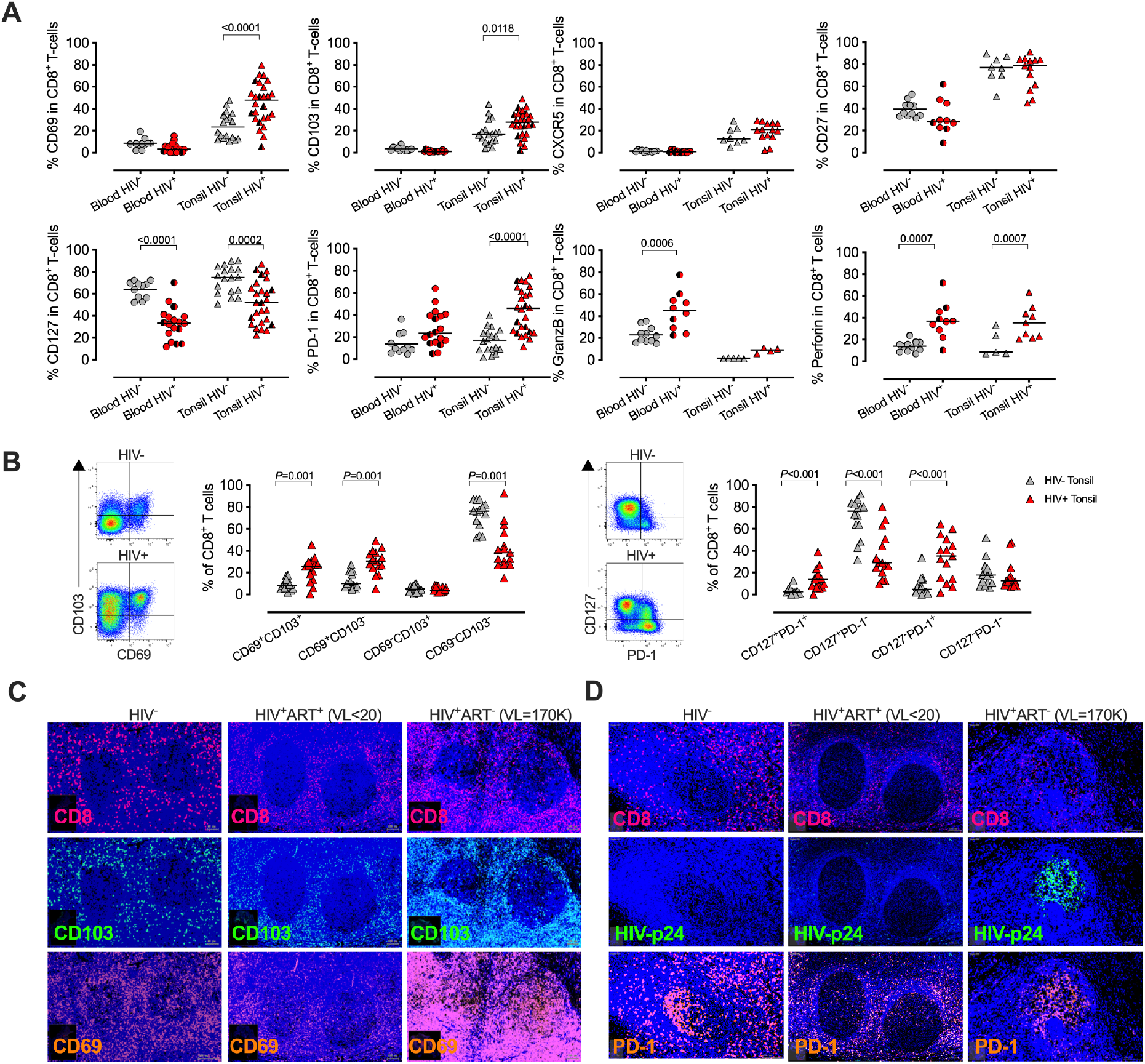
HIV infection alter CD8^+^ T-cell phenotypes in tonsils with distinct tissue locations within HIV infected and uninfected tonsils. **A**. Percentage of blood (circles) and tonsil (triangles) CD8^+^ T cells expressing CD69, CD103, CXCR5, CD27, CD127, PD-1, granzyme B and perforin markers between HIV^−^ and HIV^+^. Red data point indicates viremic (detectable HIV RNA) and red/black data point indicates suppressed (undetectable HIV RNA). P-values calculated using ordinary one-way ANOVA with horizontal bars representing median values with the level of significance indicated with p-value. **B**. Representative FACS plot of CD8^+^ T-cells from HIV-(grey triangles) and HIV^+^ (red triangles includes both ART suppressed and viremic individuals) tonsils for co-expression of CD69/CD103 and PD-1/CD127 with cumulative data shown with horizontal bars representing median values and p-values by Kruskal-wallis multiple comparisons. **C**. CD8, CD69, CD103 fluorescent immunohistochemistry of whole tonsil sections zoomed in at individual GCs for three independent donors from HIV^−^ (left), HIV^+^ART^+^ (middle) and HIV^+^ART^−^ (right) with individuals markers shown. **D**. Same as in C but for CD8, HIV-p24 and PD-1.

Next, we examined the localisation of these markers in tonsils, using fluorescence microscopy in an uninfected, viral suppressed and a viremic participant (Figure 2C). Across multiple GCs we observed that CD8^+^ cells were located outside the GC in the HIV negative and the ART suppressed individuals, but appeared to have infiltrated the GC in the HIV viremic donor. Consistent with this, CD103 was found outside GCs in the uninfected and suppressed donors, but was present within the GC and overlayed CD8^+^ staining in the viremic donor. In addition, overall staining with CD103 was increased with this tissue, consistent with the elevated CD103 expression observed by flow cytometry in PLWH. Likewise, CD69 was expressed at low levels within the GCs of HIV negative and viral suppressed tonsils, but was greatly increased both inside and outside the GC within the viremic tissue. The elevated presence of CD8^+^, CD103^+^ and CD69^+^ cells in the GCs of an HIV viremic donor was associated with high HIV-p24 detection in the same GC (Figure 2D). PD-1 was also highly expressed within these GCs, but also with HIV uninfected GCs, which, in the absence of CD8^+^ T-cells, most likely relates to T-follicular helpers cells [54, 55]. This indicates that in viremic HIV infection CD8^+^ T-cells expressing T_RM_ markers gain access to the GCs in close proximity to HIV-p24 detection. Thus, HIV infection is associated with an increase in CD8^+^ T-cells in tonsils expressing canonical T_RM_ markers, high levels of PD-1, increased perforin expression and low levels of CD127, consistent with chronic activation, increased cytolytic activity and impaired homeostatic signalling [38].

### HIV infection alters memory CD8^+^ T-cells in human tonsils

Expression of inhibitory receptors, such as PD-1, are linked to T-cell differentiation [56] and reduced functionality in the blood of PLWH [33, 37], but the role of PD-1 expression on CD8^+^ T-cells in lymph nodes remains less clear. To examine this further, we identified CD8^+^ T-cell memory subsets, based on expression of CD45RA and CCR7 [57] (Figure 3A), and found that HIV infection leads to an enrichment of effector memory and a reduction in naïve CD8^+^ T-cells in both blood and tonsil (Figure 3B). The effect of HIV on expression of CD127, PD-1 and perforin in both tonsils and blood, outlined in Figure 2, was confined to memory populations, with no significant differences observed on naïve cells (Figure 3C). The increased expression of CD69 on T_CM_ and T_EMRA_ CD8^+^ T-cells in tonsils of PLWH compared to uninfected controls observed here (Figure 3C), is consistent with recent data from HIV infected lymph nodes [38]. CD103 expression, however was unchanged across all memory subsets, suggesting the upregulation of this T_RM_ marker in PLWH was not driven by specific T-cell activation (Figure S3A).

**Figure 3:**
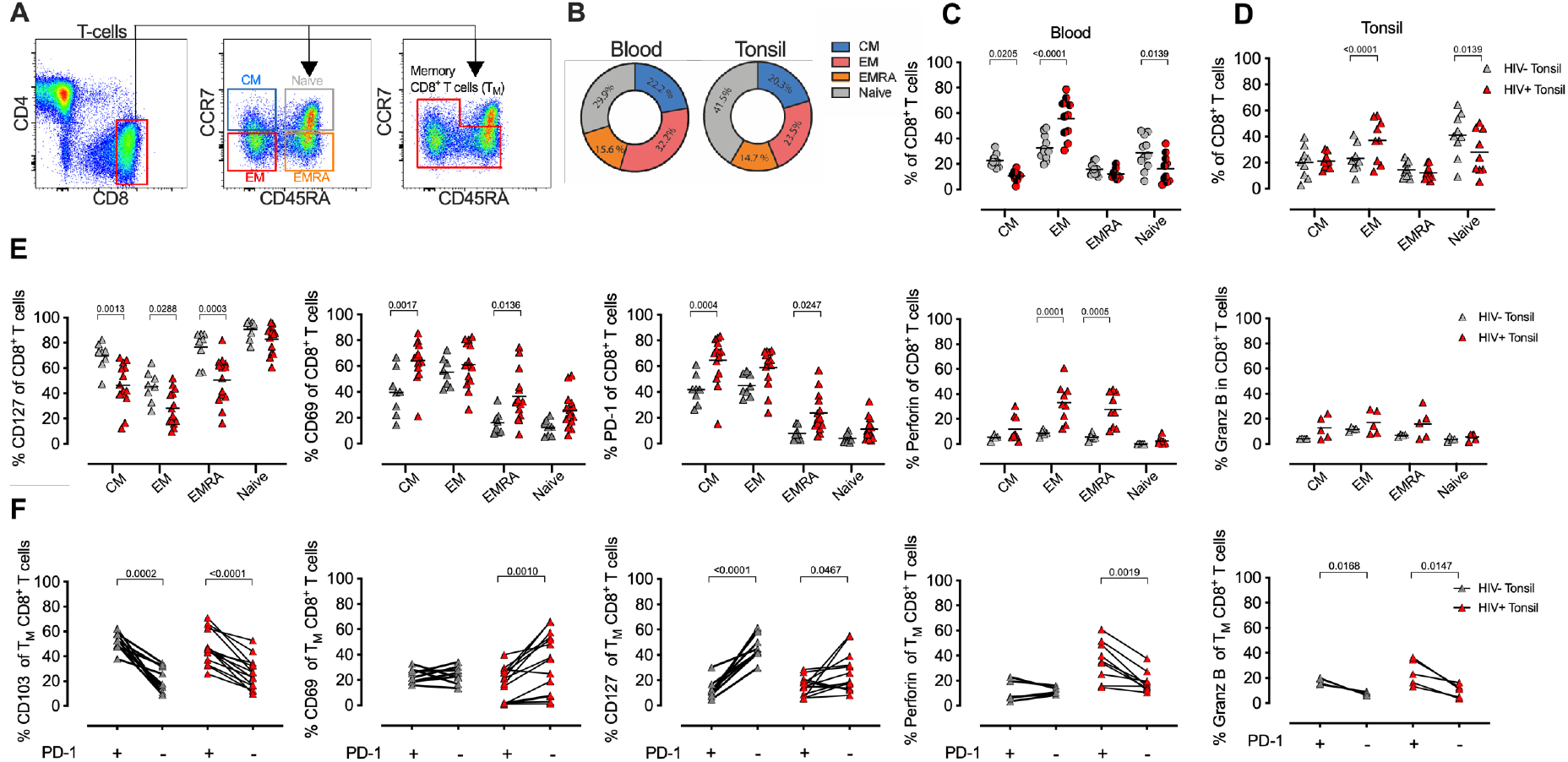
CD8^+^ T-cell memory profiling in blood and tonsils during HIV infection. **A**. Representative FACS plot of the gating strategy of CD8^+^ T-cells from tonsil cells to detect naïve, T_N_ (CCR7^+^ CD45RA^+^), central memory (T_CM_, CCR7^+^ CD45RA^−^), effector memory (T_EM_, CCR7^−^ CD45RA^−^) and T_EMRA_ (CCR7^−^ CD45RA^+^). **B**. Different memory subset distribution within blood and tonsil CD8^+^ T-cells for central memory, T_CM_ (blue), Effector memory, T_EM_ (red), transitional, T_EMRA_ (orange), T_Naive_ (grey) with cumulative memory subset distribution of CD8^+^ T cells for blood (circles, left) and tonsil (triangles, right). **C**. Distribution of blood central memory (T_CM_), transitional memory (T_EMRA_), effector memory (T_EM_) and naïve subsets within CD127, CD69, PD-1, perforin and granzyme B expressing CD8^+^ T-cells cumulative for all study participants in HIV^−^ (grey) and HIV^+^ (red). **D**. Same as in C but showing data from tonsil CD8^+^ T-cells. P-values calculated using ordinary one-way ANOVA with horizontal bars representing median values with the level of significance indicated above. **E**. The frequency of CD103, CD69, CD127, perforin, and granzyme B (Granz B) cells measured on PD-1^+^ (left) and PD-1^−^ (right) CD8^+^ T-cells from blood in HIV^+^ (red) and HIV^−^ (grey) individuals. **F**. Same as in E but showing data from tonsil CD8^+^ T-cells. P-values calculated using Paired Student’s t test. Horizontal bars represent median values.

Next, we focused on the enhanced PD-1 expression in PLWH and gated on memory subsets alone (CD8^+^ T_M_-cells) in blood and tissue and determined the co-expression of the remaining phenotypic markers (Figure 3D). In blood, although CD8^+^ T-cells expressing CD103 and CD69 are infrequent, they were generally PD1-. By contrast, PD1^+^ CD8^+^ T-cells in tonsils are generally CD103+, irrespective of HIV infection status, and CD69^−^ in PLWH. There is a strong negative association between PD-1 expression and CD127 in both the blood and tonsils of HIV uninfected participants, but this is much less pronounced in PLWH, most likely due to overall low expression levels of CD127 in these subjects. Perforin and granzyme B expressing CD8^+^ T-cells in the blood are generally PD-1^−^ (Figure S3D), consistent with the use of PD-1 as a marker of exhaustion in blood. In tonsils, however, the opposite is true, and cells expressing granzyme B in both groups and perforin in PLWH are generally PD-1^+^. Collectively, these data show that markers associated with tissue residency, immune activation and cytolytic activity are driven by memory subsets and altered by HIV infection.

### HIV-specific tonsil CD8^+^ T-cells express PD-1^high^ and CD127^low^ phenotypes compared to CMV-specific cells

Having observed phenotypic differences between blood and tonsils on bulk CD8^+^ T-cells, we next explored the role of antigen specificity in these changes and used HLA-class I tetramers to detect HIV-specific, cytomegalovirus (CMV)-specific and non-HIV/CMV specific (bulk) CD8^+^ T-cells in donor matched tonsil and blood (Figure 4A). CD69 and CD103 expression was very low or absent on both HIV and CMV-specific T-cells in the blood. In tonsils, however, the majority of HIV-specific CD8^+^ T-cells expressed both CD69 and CD103 consistent with a T_RM_-like phenotype. In contrast, although CMV-specific CD8^+^ T-cells expressed CD69 more frequently in tonsils compared to blood, this was lower than both HIV-specific and bulk CD8^+^ T-cells, and these cells almost entirely lacked CD103 expression (Figure 4B). In both compartments, PD-1 expression was significantly more frequent on HIV-specific CD8^+^ T-cells compared to CMV-specific cells. In blood, the elevated expression of PD-1 on HIV-specific CD8^+^ T-cells linked to cognate antigen stimulation [36, 37] may indicate high levels of on-going HIV antigen exposure in the tonsils of these individuals. Finally, the great majority of both HIV and CMV-specific T-cells in blood were CD127 negative, consistent with terminally differentiated CD8^+^ T-cells in circulation (Figure 4C). In tonsils, however, CMV-sepcific T-cells were largely CD127^+^, whilst HIV-specific T-cells lacked CD127 expression, consistent with the observed PD-1 expression on these cells. Thus, HIV-specific CD8^+^ T-cells in HIV-infected tonsils showed extreme phenotype changes compared to bulk and CMV-specific CD8^+^ T-cells suggesting these changes are, at least in part, driven by HIV antigenic stimulation.

**Figure 4:**
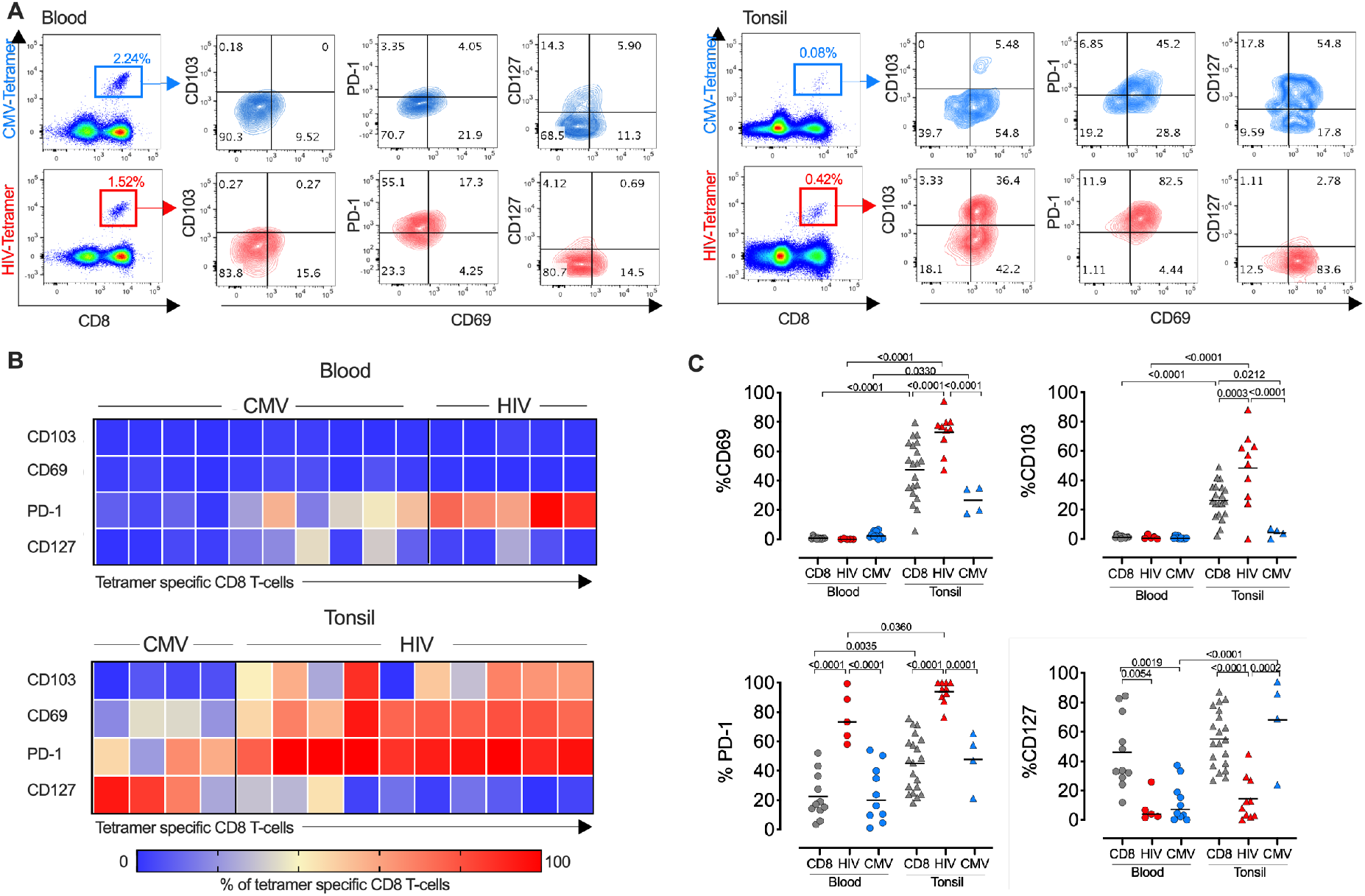
HIV-specific CD8^+^ T cells express high levels of PD-1 and T_RM_ markers in tonsils. **A**. Representative flow plots showing CMV-(top) and HIV-specific (bottom) tetramer stains of blood (left) and tonsil tissue (right) from the same participant. **B**. Heat map showing expression frequencies for the indicated markers among CD8^+^ T cells from CMV tetramer and HIV tetramer specific CD8^+^ T-cells in blood (top) and tonsil (bottom) gated CD8^+^ T-cells with frequencies for each tetramer population indicated in the bar below from blue (0%) to red (100%). **C**. Frequencies of CD69, CD103, PD-1 and CD127 from HIV-, CMV-, and non-specific (‘CD8’) CD8^+^ T-cells within blood (left) and tonsil (right) tissue. P-values calculated using ordinary one-way ANOVA with horizontal bars representing median values with the level of significance indicated above.

### Single-cell transcriptional profiling identifies distinct clusters of HIV-specific and activated PD-1^high^ tonsil CD8^+^ T-cells linked to exhaustion signatures

To further characterise how HIV infection impacts HIV-specific CD8^+^ T-cells in tonsils, we applied an optimized SMART-seq2 protocol to perform scRNAseq on single-cell tetramer sorted HIV- and CMV-specific CD8^+^ T-cells as well as bulk CD8^+^ T-cells (non-specific CD8^+^ Tet^−^), from tonsils of participants with chronic untreated HIV infection (Table S2, Figure 5A). We first created a cells-by-genes expression matrix, performed variable gene selection and unsupervised clustering using the dimensionally reduction by tSNE with graph-based clustering and identified four distinct CD8^+^ T-cell clusters (Figure 5B). Although the relative frequency in each cluster varied between donor, we found cells from each donor represented in all of the 4 clusters. Furthermore, all of the 7 different tetramer specific and non-specific CD8^+^ T-cells were represented in each cluster, suggesting they describe differences in underlying cellular states rather than donor and/or antigen specific effects. Differential gene expression analysis among these four clusters revealed notable molecular differences (Figure 5C and Table S3). Genes in cluster 0 (blue) included *JUN, JUNB, FOS, FOSB* and *ATF*, which together form the AP-1 transcription factor, and the MAPK signalling genes *DUSP1* and *DUSP2*, both of which control cellular differentiation and proliferation during viral infection [58] along with *EGR2*, which is also expressed by cluster 0 [59, 60]. Together with the expression of *TNFSF9*, involved in lymphocyte activation and *IFNG*, this transcriptional signature suggests thaT-cells in cluster 0 are highly activated and proliferating. Cluster 1 (orange) marks genes involved in activation – *HLA-E, CFL1, STAT3, CD97, LMNA* and along with the transcription factor *KLF6*. Cluster 2 contains mostly ribosomal genes and potentially indicates increased activity of protein machinery, as observed in the lymph nodes of elite controllers [43]. Cluster 3 contains *IL17RD*, involved in the TH17 response, and *CCL22*, interacting with CCR4, along with the TNF-induced protein 8 encoded by *TNFAIP8L1* and *AXL* involved in tyrosine kinase signalling (Figure 5C).

**Figure 5:**
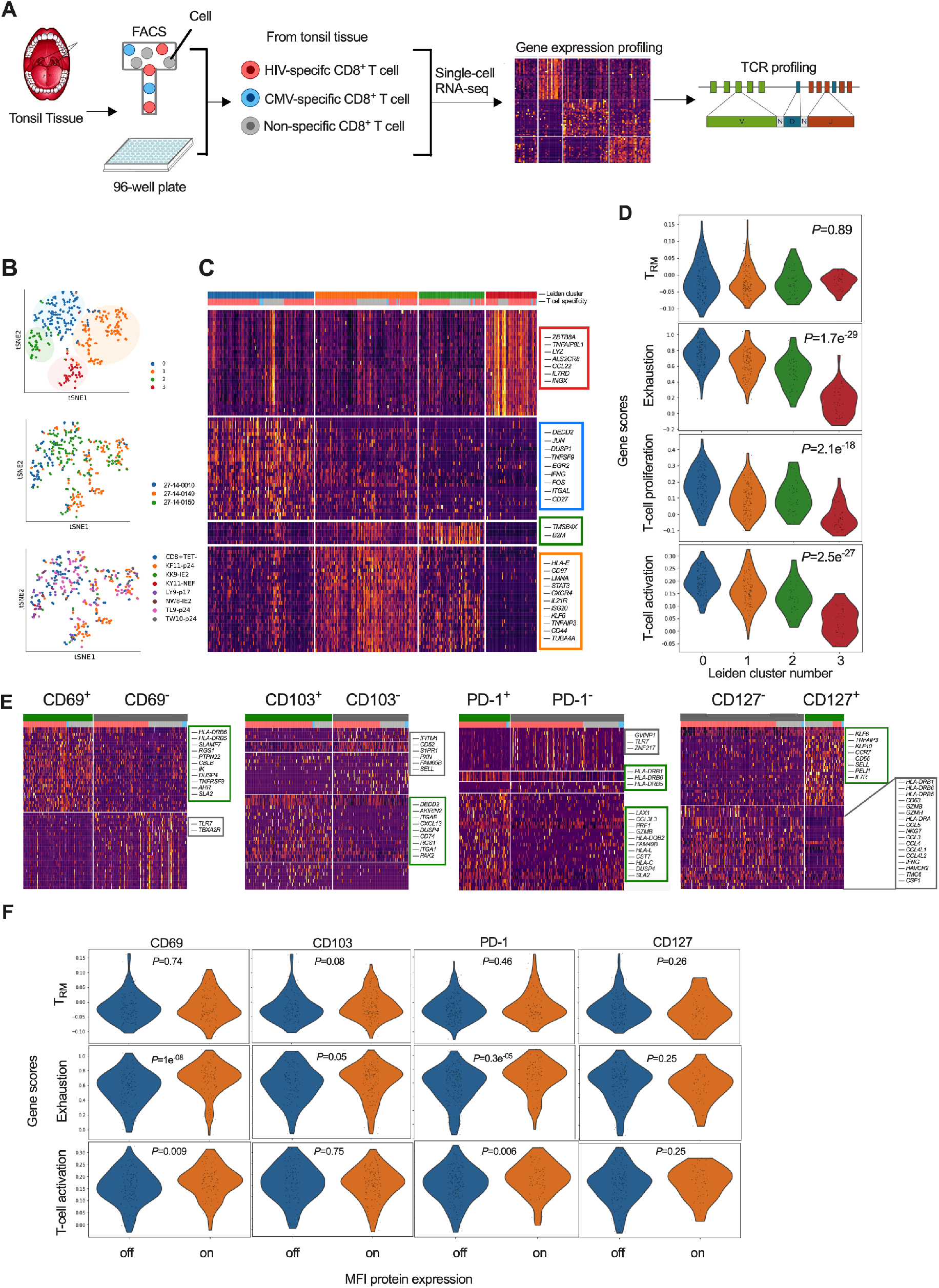
Single cell transcriptional profiling of CD8^+^ T-cells within HIV infected tonsils reveal distinct cellular subsets grouped by T-cell exhaustion gene profiles. **A**. Workflow of single-cell RNA sequencing from HIV infected tonsil tissue isolated CD8^+^ T-cells pre-sorted on HIV-, CMV- and ‘non-specific’ CD8^+^ T-cells from HIV infected participants indicated in Table S2. **B**. Dimensionality reduction using tSNE on scRNA-Seq cells coloured by Louvain cluster (top), participant ID (middle), and tetramer specificity (bottom). **C**. Heatmap of z-scored gene expression of top differentially expressed genes (t-test) between Louvain clusters from scRNA-seq data with cells grouped by Louvain cluster, genes grouped by hierarchical clustering (full gene lists in supplemental Table S3). **D**. Gene set enrichment scores for each of the 4 Louvain clusters (0-3) shown for ‘ T_RM_’ [38], ‘Exhaustion’ [61], ‘proliferation’ and ‘activation’ [62, 63] published gene lists. **E**. Heatmaps of z-scored gene expression of top differentially expressed genes (t-test) between single cells with high and low normalized MFI values of CD69, CD103, PD-1 and CD127. Selected genes labelled in plot, full gene lists in Table S4-S5. **F**. Gene lists from (E) scored against the published gene lists as indicated in D.

To ascribe functional states to these 4 gene clusters (‘Leiden clusters 0-3’) we scored them against gene-sets derived from: i) tissue resident memory CD8^+^ T-cells (T_RM_) in human lymph nodes [38] (Table S2); ii) exhausted T-cells [61] (Table S13); and, iii) T-cell proliferation and iv) T-cell activation [62, 63]. We found no difference in T_RM_ scores (P=0.89), but a strong difference in exhaustion, proliferation and activation profiles (P<0.0001), which were highest for cluster 0 followed by cluster 1 and 2. Cluster 3 scored low in all gene scores, except T_RM_, indicating more quiescent cellular states despite majority of these cells are HIV-specific and T_RM_-like (Figure 5D). As shown in the heatmap, HIV-specific (highlighted in red), CMV (blue) and bulk (grey) CD8^+^ T-cells were found in all 4 clusters. Thus, HIV-specific CD8^+^ T cells do not exclusively express an exhausted or activated phenotype with no clear evidence of differential T_RM_ gene set enrichment between these different antigen specificities.

### CD69 and PD-1 expression on HIV-specific CD8^+^ T-cells are linked to activated immune exhaustion signatures

Having observed differences in CD103 and PD-1 expression by flow cytometry between CMV and HIV-specific CD8^+^ T-cells, we next directly compared gene expression levels (mRNA) with matched surface protein expression (MFI) on the index sorted cells. Overall, we found higher protein expression of CD69, CD103 and PD-1, and lower CD127 expression on HIV-specific compared to CMV-specific CD8^+^ T-cells, consistent with bulk analysis above (see Figure 4) (Figure S3A). This was consistent with gene expression for *PDCD1* (PD-1) and *IL7R* (CD127), but not for *CD69* or *CD103*. Accordingly, a direct correlation was found between MFI and mRNA abundance for IL7R (CD127) but not for CD69, CD103 and PD-1 (Figure S4B) potentially due to shorter mRNA half-lifes compared to protein stability [64]. To test whether differences in protein expression of these markers were linked to specific gene signatures, despite inconsistent detection of the genes themselves, we applied MFI clustering based on our protein index markers to generate ‘on-off’ signatures for each protein marker (Figure S3C), and used that to directly compare genes associated with protein expression (Figure 5E). Using this approach, cell surface expression of CD69 was found to be associated with *HLA-DRB6, HLA-DRB5, SLAMF7* and *TNFRSF9* (4-1BB/CD137), known to regulate CD8^+^ T-cell clonal expansion and associated with recent TCR-triggered activation [65] (Table S4) suggesting CD69 expression may be associated more with T-cell activation in tonsils rather than tissue residency. CD103 expressing cells, by contrast, were associated with CD8^+^ T_RM_ markers, such as the adhesion markers *ITGA1* (CD49a), the chemokine receptor *CXCL13*, and *RGS1*, a G-protein-signalling gene involved in modulating T-cell trafficking (Table S5) [66]. In contrast, CD103^−^ cells expressed genes associated with tissue egress, such as *S1PR1* and its associated transcription factor *KLF2* along with *SELL* (CD62L), suggesting that these cells can re-circulate via lymph and peripheral blood [38, 49, 67]. PD-1 expressing CD8^+^ T-cells where almost exclusively HIV-specific and expressed genes linked to cytotoxicity (i.e., *PRF1, GZMB, CCL3L3, CST7*) and activation (*HLA-DR, HLA-DQ, HLA-L, HLA-C*) (Table S6). Finally and in line with flow cytometry data on bulk CD8^+^ T-cells, CD127 expression marked naïve like cells, identified through expression of *CCR7, SELL* and *IL7R*, and was dominated by non-specific CD8^+^ T-cells in constrast to HIV-specific CD8^+^ T-cells that were mostly CD127^−^ and associated with upregulation of chemokine ligands (e.g., *CCL4L2, CCL4L1, CCL3, CCL4, CCL5*), IL-12-induced effector molecules (*IFNG, GZMH, GZMB, NKG7*) and genes involved in T-cell activation (*HLA-DRA, HLA-DRB1, HLA-DRB6, HLA-DRB5, CD63*) (Table S7), which is consistent with the downregulation of CD127 on activated HIV-specific CD8^+^ T-cells (see Figure 4D).

Next, we scored these gene sets against the T_RM_, exhaustion and T-cell activation gene lists used above (see Figure 5D). No significant differences in their T_RM_ scores was observed for any of the proteins measured (Figure 5F). However, the T_RM_ score was higher in CD103^+^ cells, and this difference approached significance (P=0.08), consistent with the upregulation of T_RM_ associated genes highlighted above. Analysis of the exhaustion gene list showed both PD-1 and CD69 protein expression were strongly associated with CD8^+^ T-cell exhaustion (P<0.0001), but not CD127 expression (P=0.24) (Figure 5F). Finally, T-cell activation scored highest in CD69 and PD-1 expressing cells, but with reduced CD127 in HIV-specific cells compared to non-specific cells, suggesting that CD69 may act as an activation marker within HIV-infected tonsil CD8^+^ T-cells (see Figure 4). These data suggest that the phenotypic differences observed on HIV-specific CD8^+^ T-cells in tonsils of viremic individuals are associated with the specific activation and exhaustion profiles potentially linked to persistent HIV antigen in these tonsils, although not all HIV-specific cells expressed this phenotype (see Figure 5D).

### HIV-specific CD8^+^ T-cells from ‘controller’ tonsil are non-cytolytic compared to viremics

Finally, one of our HIV infected tissue donors maintained undetectable plasma viral load in the absence of ARV detected drugs in plasma, subsequently referred to as “controller” (see Table S2). This donor possesses protective HLA class I alleles that present two HIV epitopes KF11-p24 and TW10-p24 CD8^+^ T-cells associated with viral control [20, 68, 69] and offered the opportunity to determine the signatures of HIV-specific CD8^+^ T-cells isolated from tonsils where HIV antigen burden is low. First, we confirmed the low level of antigenemia in the controller by immunohistochemistry, and observed extremely low levels of HIV-p24 within the GCs compared to tonsils from viremic donors highlighting the GCs as HIV sanctuary sites (Figure 6A). Next, we performed direct comparison of scRNAseq profiles of HIV-specific CD8^+^ T-cells from the controller and the two viremic individuals analysed (Figure 6B). This identified genes associated with cytolytic activity (*GZMA, GZMB, GZMH, GZMK*), interferon stimulating gene 20 (*ISG20*) and its downstream *STAT3* signalling in HIV-specific CD8^+^ T-cells from viremics compared to the controller (Figure 6B, Table S12). In contrast, cells from the viral controller expressed transcripts involved in the AP-1 transcription factor complex controlling cellular differentiation *TNFSF9, TNFS14, TNFS15* and *JUN, JUNB* (Figure 6B). Overall, this is consistent with a distinct transcriptional profile observed in lymph node CD8^+^ T-cells from elite controllers, which notably lacked cytolytic activity [43]. Pathways analysis of 1,622 significantly DEGs revealed distinct functional differences between cells from viremic and controller tonsils (Figure 6C). In particular, we found that viremic donors express elevated pathways that included IL-15 signalling, Th1 pathways, extravasation signalling, T-cell exhaustion signalling, Granzyme B signalling and increased metabolic activity, such as oxidative phosphorylation, mTOR and EIF2 signalling (Figure 6C, Table S13). Upstream driver analysis predicted activating cytokines involved in type I T-cell responses, including IL-15, IL-2, IFN-γ and TNF-↵ (Figure 6D, Table S9). Matched protein expression of PD-1 and CXCR5 on single cells from tonsils obtained from non-controller and controllers clearly indicates reduced expression of PD-1 on HIV-specific CD8^+^ T-cells from the controller, in addition to CXCR5 (Figure 6E).

**Figure 6:**
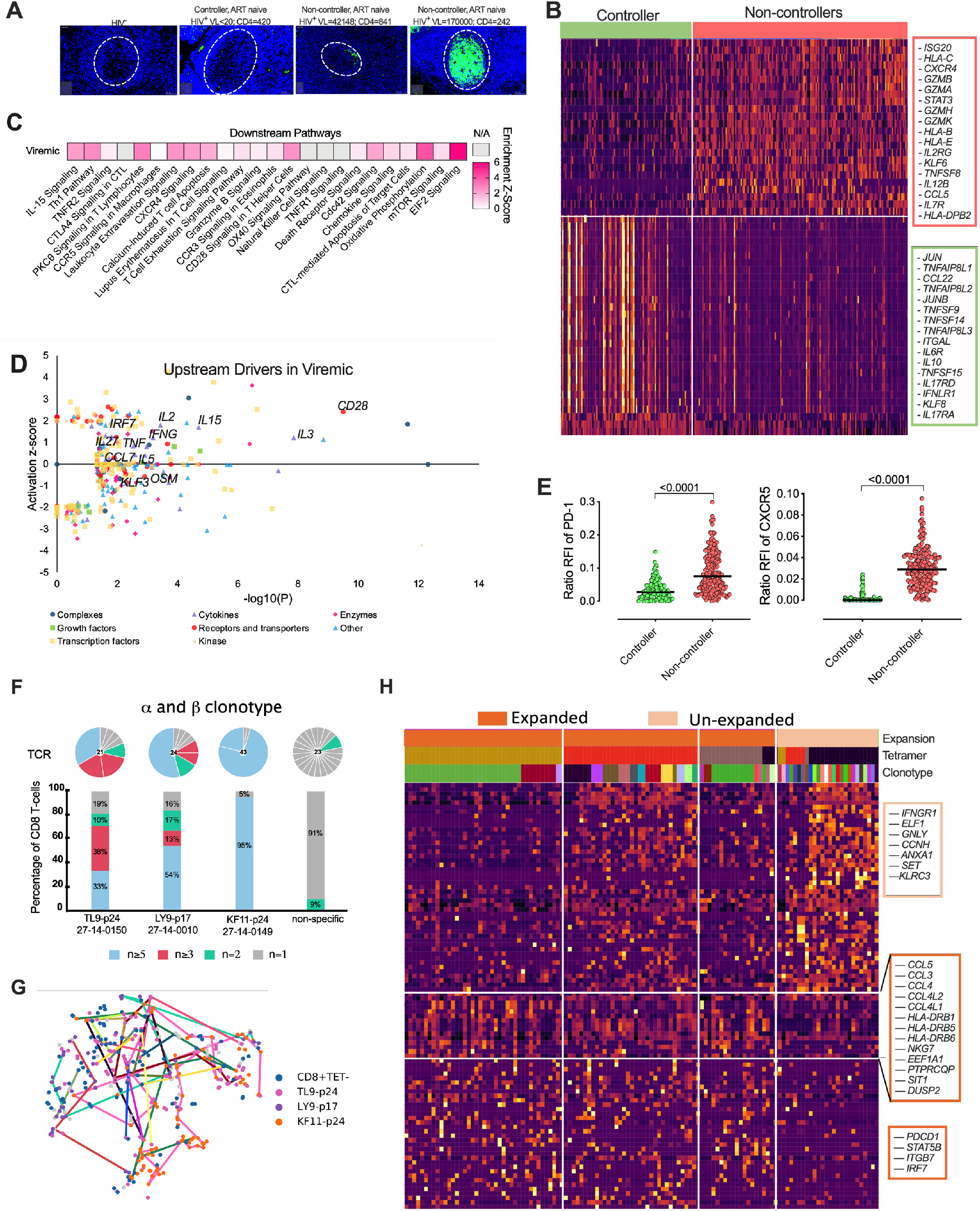
Natural HIV ‘controllers’ display reduced HIV-specific CD8^+^ T-cell cytolytic transcriptional signatures and PD-1 expression compared to viremic ‘non-controllers’ and linked to *α-β* TCR clonotypes. **A**. Fluorescent immunohistochemistry of HIV-24 protein (green) for 4 donors of HIV^−^, HIV^+^ ‘controller’ and two HIV^+^ART^−^ ‘non-controllers’ with plasma viral load and absolute blood CD4 counts listed above. **B**. Heatmap showing differentially expression of 300 featured genes with selectively specifically expressed genes marked (right) among 1.622 differentially genes (Table S12). **C**. Selected canonical pathways by ingenuity pathway analyzer. **D**. Upstream drivers of pathways significant by ingenuity pathway analysis (IPA) of DEGs. For directionally annotated pathways, a Z-score is calculated to represent up- or downregulation of the driver or pathway. If a driver or pathway is not directionally annotated in IPA, or there are not enough genes in the list to calculate a Z-score, N/A is reported. See Table S9 for the full IPA results. E. Relative fluorescent intensity (RFI) for PD-1 and CXCR5 expression for each HIV-specific CD8 T-cell comparing ‘controller’ and ‘non-controller’ participants. **E**. Relative fluorescent intensity (RFI) for PD-1 and CXCR5 expression for each HIV-specific CD8 T-cell comparing ‘controller’ and ‘non-controller’ participants. **F**.The TCR distribution of KF11-p24, TL9-p24 and LY9-p17 from 27-14-0149, 27-14-0150 and 27-14-0010, respectively (see Table S2) with unique (n = 1), duplicated (n = 2), triplet (n 3), and clonal (n 5) with bars colored in grey, green, pink and blue representing the fraction of cells belonging to groups of clonotypes with either 1, 2, 3-4 or more than five clonotypes, respectively. Pie chart above each bar illustrate the composition of every individual *α-β*-TCR. **G**. tSNE projection of Smart-Seq2-profiled flow-sorted CD8^+^ T cells on the lovain clusters (see Figure 5B) with colored lines connecting cells sharing the same CDR3 sequence. **H**. Heatmap of z-scored gene expression of top differentially expressed genes (t-test) between Louvain clusters from scRNA-seq data. Cells grouped by expansion of clonotype, genes grouped by hierarchical clustering. Full gene lists in Table S10.

Next, we tested the clonal distribution of a subset of these HIV-specific and bulk non-HIV-specific CD8^+^ T-cells from controller and non-controllers (see Figure 5A) and found clear *α-β* matched clonotype expansion of HIV-specific cells as expected (Figure 6F), and by ↵ and /3 chains separately (Figure S5B,C). We linked each HIV-specific paired *α-β* TCR clonotype to their transcriptome (see Figure 5B) and found that enriched clonotypes were represented across 2 or more of the different transcriptional clusters (Figure 6G and see Figure 5B), indicating that the TCR:antigen interaction does not determine transcriptional state textitper se. However, gene expression analysis between enriched vs unenriched clonotypes revealed a signature of enriched *α-β* clonotypes, including upregulated HLA class II genes, suggesting activation (*HLA-DRB1/DRB5/DRB6*), combined with chemokine and chemokine ligands (*CCL3/4/5, CCL4L1/2*) involved in T-cell trafficking and homing (Figure 6H). In addition, this gene signature was enriched for *PDCD1*, encoding PD-1, consistent with the hypothesis that PD-1 expression is driven by T-cell activation and expansion. Enriched clones also expressed higher levels of integrin *ITGB7* and *IRF7* involved in IFN-↵ induction in lymphoid tissue, and *STAT5B* signalling linked to TCR signalling. By contrast, unexpanded cells, defined by unique clonotypes, expressed lower levels of these genes associated with T-cell activation, but retained expression of genes associated with CD8^+^ T-cell function (*IFNGR1, GNLY, SET, ANZA1, ELF1* and *KLRC3*), in addition to multiple housekeeping genes (Figure 6H, Figure S5D, Table S9-S11).

These data support the hypothesis that the activated and exhausted phenotypes observed in HIV-specific CD8^+^ T-cells in tonsils are driven by antigen specific activation, or its inflammatory sequelae, because they are reduced in the absence of antigenic stimulation in controllers. Consistently, expanded clonotype transcriptomes are also associated with signatures of activation, trafficking and exhaustion. These data supports the potential importance of non-cytolytic CD8^+^ T-cell activity in controlling viral replication within lymphoid tissues from natural HIV controllers [43].

## Discussion

Over recent years, in-depth studies have advanced our understanding of T_RM_ in human organs and have highlighted the distinction between peripheral blood CD8^+^ T-cells and CD8^+^ T_RM_ cells in particular [70, 71]. The past decade has also established that a major HIV viral reservoir, responsible for viral rebound after cessation of antiretroviral treatment, is located in mucosal and lymphoid tissue sites [72–74]. However, little is known about antiviral HIV-specific CD8^+^ T-cells located in close proximity to HIV infected cells in these tissue sites, which is important if they are to be leveraged to help purge the HIV reservoir.

In this study, we made an in depth analysis of CD8^+^ T_RM_-like cells in HIV infected tonsils by applying transcriptional, functional, clonotype and phenotypic characterisation of bulk and HIV-specific CD8^+^ T-cells combined with *in situ* HIV-p24 detection. We found that HIV infection has a dramatic impact on the expression of CD8^+^ T_RM_ markers in tonsils and that the majority of HIV-specific CD8^+^ T-cells express very high levels of PD-1, CD69 and CD103 and low levels of CD127. These differences were much greater in HIV-specific CD8^+^ T-cells compared to either CMV specific or bulk CD8^+^ T-cells in the same tonsils. Our single-cell transcriptional profiling revealed subclustering of CD8^+^ T-cells in tonsils and indicates that the upregulation of CD69, which serves as both a marker of T_RM_ and activation, and PD-1 on these cells is linked to transcriptional signatures of activation, differentiation, interferon stimulating genes and exhaustion. Moreover, CD8^+^ T-cell ancestry analysis suggest that these exhaustion signatures are enriched in expanded HIV-specific clonotypes, suggestive of on-going proliferation. Finally, PD-1 and CD69 expression was much lower on HIV-specific CD8^+^ T-cells isolated from the tonsils of a natural controller. In contrast to the non-controllers, these cells displayed enhanced expression of non-cytolytic T-cell genes, associated with immune control on elite controllers, and lacked the signatures of activation and immune exhaustion.

Our findings also indicate that the subset composition and phenotype of peripheral blood CD8^+^ T-cells does not reflect that of tonsil tissue, highlighting the need to study relevant tissue sites. Single-cell transcriptional profiling comparing tonsil and blood defined core CD8^+^ T_RM_ signatures characterised by lower cytolytic programs, whereas blood CD8^+^ T-cells showed consistent re-circulating profiles. This is inline with other recently published data showing that cytolytic CD8^+^ T-cells are restricted to the intravascular circulation [49], and questions the relationship between T-cell functionality measured in the blood and what manifests with lymphoid structures, such as the tonsil.

Both CD8^+^ T-cells from blood and tonsil showed a predominantly memory phenotype that in tonsil tissue were enriched for CD69, which is consistent with expression of transcription factors involved in preventing tissue egress. We found increased CD69 expression on CD8^+^ T_EM_ from HIV infected tonsils that was almost double that of CD8^+^ T_EM_ cells from HIV uninfected tonsil and consistent with observations in lymph nodes [38]. In contrast to conventional lymph nodes, palentine tonsils belong to the oral mucosal tissue and possess crypts and an epithelial barrier expressing E-cadherin, a ligand for CD103 involved in tissue retention of CD8^+^ T_RM_ [17, 75], which we found upregulated by HIV infection and co-expressed with CD69. Interestingly, approximately half of the CD69 expressing CD8^+^ T-cell population in HIV infected tonsils were CD103^+^, which is distinct to that found in LNs and highlights the existence of compartment specific differences between these different secondary lymphoid organs [38]. The upregulation of CD69 and CD103 may suggest that HIV infection increases the retention of CD8^+^ T_RM_ cells in tonsils, which could be well placed to participate in a durable response to ongoing viral replication in lymphoid tissue.

Elevated CD8^+^ T_RM_ marker expression is also consistent with our observations of strong upregulation of both PD-1 and cytolytic markers on bulk and memory CD8^+^ T-cells. Thus, the increased frequency of CD8^+^ T_RM_ cells in HIV-infected tonsils suggests that these cells are continuously engaged in antiviral activities at the site of the viral reservoir. Interestingly, we found the majority of CD8^+^ T-cells located outside the GCs, but that GCs contained the vast majority of the HIV reservoir suggesting that CD8^+^ T-cells fail to accumulate in large numbers in B-cell follicles [76] and adds to the limited knowledge of the location of HIV-specific CD8^+^ T-cells *in situ* [39].

When we performed a peptide-MHC tetramer-driven analysis of antigen specific CD8^+^ T-cells, we consistently found increased expression of T_RM_ markers and PD-1 expression on HIV-specific compared to CMV specific CD8^+^ T-cells in tonsils. In particular, the HIV-specific CD8^+^ T-cells were enriched in the distinct T_RM_ and PD-1^high^/CD127^low^ phenotype compared to the matched bulk CD8^+^ T-cells and further support that these cells are retained in HIV infected tonsils where they are involved in the antiviral response. This is consistent with our unsupervised clustering of HIV-specific T_RM_-like cells by single-cell transcriptomes that revealed that distinct subsets exist related to exhaustion, activation and proliferation. We also found distinct transcriptional gene sets by cell surface marker separation alone. For example, CD69 and PD-1 expression appeared to be associated with gene sets of exhaustion and activation in contrast to CD127 expression that was enriched in the non-HIV-specific cells and consistent with a naïve-like and re-circulating transcriptional profile [49]. This is also consistent with the *SELL, KLF2* and *S1PR1* transcripts within the CD103^−^CD8^+^ T-cells. Thus, re-circulating signatures were enriched in the CD127^+^ and CD103^−^CD8^+^ T-cells that also contains the majority of non-HIV-specific CD8^+^ T-cells in this analysis and further supporting that HIV-specific CD8^+^ T-cells are tissue resident [38, 43].

The inclusion of tonsil tissue from a natural HIV controller allowed us to compare the HIV-specific CD8^+^ T-cells to those purified from viremic ART naïve individuals. This is to our knowledge the second observation in human tissue [43] and the first in tonsils. We found distinct signatures showing that CD8^+^ T_RM_- like cells within HIV ‘controlled’ tonsils lacked transcripts encoding cytolytic effector molecules, such as *GZMA, GZMB, GZMH* and *GZMK*, while tonsil CD8^+^ T-cells from viremic individuals was characterized by upregulated pathways involved in IL-15 signalling, granzyme B signalling, T-cell apoptosis/exhaustion and oxidative phosphorylation. Thus, the active metabolisms in these cells support our hypothesis above that increased antigen load in viremic compared to HIV controlled tonsils is linked to CD8^+^ T-cell activation and exhaustion. Moreover, the lower cytolytic profile from HIV controlled tonsil CD8^+^ T-cells is consistent with a recent study in HIV infected lymph nodes from elite controllers and further support the hypothesis of non-cytolytic features of CD8^+^ T_RM_ in elite controllers [43]. Whether HIV-specific CD8^+^ T-cells within lymphoid tissue from viremic donors can be reverted into low exhaustive and low cytoytic elite-like controller function by PD-1/PD-L1 inhibition [77, 78] remains to be seen [79]. However, elite control may not solely be explained by their cellular immune responses, but also linked to distinct host genome viral integration sites [80]

In summary, we show that HIV infection enrich for CD8^+^ T_RM_ – like phenotypes located outside the GCs in human tonsils collected from PLWH in HIV endemic areas in South Africa and that the HIV-specific tonsil CD8^+^ T-cells match a non-vascular transcriptional program [49] with high levels of PD-1 and exhaustive transcriptional signatures linked to expanded clonotypes. This, combined with low PD-1 and cytolytic profiles in viral ‘controllers’, suggests that immunotherapy should be targeted to include CD8^+^ T_RM_ cells relevant for HIV cure or remission strategies recently explored in animal models [81, 82].

## Material and Methods

### Study participants

Human tonsils tissue samples and peripheral blood were collected from individuals being HIV^−^ (n=20), HIV^+^ chronic and naïve to ART (n=8), HIV^+^ chronic on ART (n=15). Tonsil tissue were obtained from patients undergoing routine tonsillectomy at Stanger Hospital, KwaDukuza and Addington Hospital in Durban, KwaZulu-Natal.

Frozen peripheral blood samples HIV- (n=10) were obtained from the Females Rising through Empowerment Support and Health (FRESH) cohort from Umlazi, Durban (Ndlovu et al., 2015). Frozen peripheral blood samples HIV+ naïve to ART (n=5) and on ART (n=5) were obtained from the ‘GATEWAY’ based at the Prince Mshiyeni Memorial Hospital, Umlazi, Durban.

All participants provided informed consent and the study was approved by the respective institutional review boards including the Biomedical Research Ethics Committee of the University of Kwazulu-Natal in Durban, South Africa.

### Sample processing-blood and tonsils

Blood samples were processed from fresh peripheral blood mononuclear cells (PBMCs) purified using Ficoll separation. Samples were thawed in DNase-containing (25 units ml-1) R10 (Sigma-Aldrich) at 37°C. Cells were rinsed at rested at 37°C for a minimum of 1h before undergoing red-blood-cell lysis by 5-10 ml RBC lysis solution (Qiagen) for 20 min at room temperature. Cells were then stained with the appropriate antibody panel described in Flow cytometry.

Tonsil samples were processed from fresh tissue immediately after surgery. Resected tissues were washed with cold HBSS (sigma-Aldrich) and dissected into smaller pieces. Tissues were rinsed again and resuspended in 10 ml R10, containing DNase (1 μl ml-1) and collagenase (4 μl ml-1), and disassociated in a GentleMACS dissociator (Miltenyi Biotec). Cells were rested in a shaking incubator at 37°C for 30 min and then further processed in the GentleMACS dissociator. After further resting (30 min at 37°C) and washing steps, cells were strained through a 70-μm cell strainer and washed on the final time. Cells were lysed using 5-10 ml RBC lysis buffer (Qiagen) and stained for flow cytometry analysis. In the relevant experiments, tetramer was added for 30 minutes at room temperature after resting. For identifying HIV and CMV-specific CD8^+^ T-cells.

### Flow Cytometry

For FACS analysis, different antibody panels for phenotype and intracellular cytokine staining (ICS) were used. A complete list of antibodies used with identifier and source information can be found in the supplementary table 14.

Cryopreserved PBMC and TMC were thawed and rested in RPMI-1640 media (20% fetal calf serum (FCS), 1% penicillin/streptomycin, 1% L-glutamine) for 2 hours at 37°C incubator. In the relevant experiments, tetramer was added for 30 minutes at room temperature after resting for identifying HIV and CMV-specific CD8^+^ T-cells. All Samples were further surface stained including a near-infrared live/dead cell viability cell staining kit (Invitrogen) at room temperature for 20 min. Cells were then washed with PBS and acquired using the BD FACS ARIA FUSION (BD Bioscience). For experiments involving ICS, the cells were stimulated with SEB, in the presence of Golgiplug and Golgistop (BD Biosciences) for 6 hours in 37°C incubator. Cells were stained with fluorochrome-conjugated monoclonal antibodies and subsequently fixed, permeabilized and stained by BD Cytofix/cytoperm kit (BD Biosciences). Peptide-MHC tetramers were acquired from ImmunAware, Copenhagen, Denmark and stained during cell surface antibody labelling. Blocking with 20% goat serum for 20 min was done prior to intracellular antibody staining. After staining cells were washed and fixed in 2% paraformaldehyde before acquisition on a 4 laser, 17 parameter BD FACSAria Fusion flow cytometer. Data were analyzed with FlowJo software (version 10.4.2, TreeStar).

### Fluorescent immunohistochemistry

Multiplex fluorescent immunohistochemistry experiment was performed using the Opal™ 4-Color Manual IHC kit (PerkinElmer) according to the manufacturer’s instructions. Briefly, tonsil tissue samples fixed with 4% formalin for a minimum of 48 hours were paraffin-embedded. Exactly 4μm sections were cut, deparaffinized and stained with the following unlabelled primary antibodies: CD8 (clone: C8/144B, Dako), CD4 (clone: 4B12, Dako), p24 (clone: Kal-1, Dako), CXCR5 (clone: MU5UBEE, Thermofisher Scientific), Granzyme B (clone: 23 H8L20, Thermofisher Scientific), CD103 (clone: NBP1-88142, Novus Biologicals) and CD69 (clone: 15B5G2, Novus Biologicals). Opal fluorophores: FITC (Opal520) was used for p24 and CD103; Texas-Red (Opal570) was used for CD8 and CD4; then Cy5 (Opal690) was used for CD8, CXCR5, Granzyme-B and CD69 signal generation in the different IHC experiments performed. DAPI was used as the nuclear counterstain. Images were acquired on a Zeiss Axio Observer Z1 inverted microscope (Olympus) and analysed with TissueFAXS imaging and TissueQuest quantitation software (TissueGnostics).

### Single-cell RNA-seq using Seq-Well S3

After obtaining single-cell suspension from fresh biopsies, we used the Seq-Well S3 platform. Full methods on implementation of this platform is described [46, 47]. Briefly 15,000 cells in 200 mL RPMI + 10% FBS were loaded onto one PDMS array preloaded with barcoded mRNA capture beads (ChemGenes) and settled by gravity into distinct wells. The loaded arrays were washed with PBS and sealed using a polycarbonate membrane with a pore size of 0.01 μm, which allows for exchange of buffers but retains biological molecules within each nanowell. Arrays were sealed in a dry 37°C oven for 40 min and submerged in a lysis buffer containing guanidium thiocyanate (Sigma), EDTA, 1% beta-mercaptoethanol and sarkosyl (Sigma) for 20 min at room temperature. Arrays were transferred to hybridization buffer containing NaCl (Fisher Scientific) and supplemented with 8% (v/v) polyethylene glycol (PEG, sigma) and agitated for 40 min at room temperature, mRNA capture beads with mRNA hybridized were collected from each Seq-Well array, and beads were resuspended in a master mix for reverse transcription containing Maxima H Minus Reverse Transcriptase (ThermoFisher EP0753) and buffer, dNTPs, RNase inhibitor, a 50 template switch oligonucleotide, and PEG for 30 min at room temperature, and overnight at 52°C with end-over-end rotation. Exonuclease I treatment (New England Biolabs M0293L) was used to remove excess primers. After exonuclease digestion, bead-associated cDNA denatured for 5 min in 0.2 mM NaOH with end over end rotation. Next, beads were washed with TE + 0.01% tween-20, and second strand synthesis was carried out by resuspending beads in a master mix containing Klenow Fragment (NEB), dNTPs, PEG, and the dN-SMRT oligonucleotide to enable random priming off of the beads. PCR amplification were carried out using KAPA HiFi PCR Mastermix (Kapa Biosystems KK2602) with 2.00 beads per 50 μL reaction volume. Post-whole transcriptome amplification, libraries were then pooled in sets of six (12,000 beads) and purified using Agencourt AMPure XP SPRI beads (Beckman Coulter, A63881) by a 0.6x volume ratio, followed by a 0.8x. Libraries size was analysed using an Agilent Tapestation hsD5000 kit (Agilent Genomics) with an expected peak at 1000 bp and absence of smaller primer peaks. Libraries were quantified sing Qubit High-Sensitivity DNA kit and preparation kit and libraries were constructed using Nextera XT DNA tagmentation (Illumina FC-131-1096) using 800 pg of pooled cDNA library as input using index primers with format as in Gierahn et al [46]. Amplified final libraries were purified twice with AMpure XP SPRI beads as before, with a volume ratio of 0.6x followed by 0.8x yielding library sizes with an average distribution of 650-750 pb. Libraries were pooled and sequenced together using a Illumina NovaSeq 6000 S2 Reagent kit v1.5 (100 cycles) using a paired end read structure with custom read 1 primer: read 1: 20 bases with a 12 bases cell barcode and 8 bases unique molecular identifier (UMI). Read 2: 82 bases of transcript information, index 1 and index 2: 8 bases.

### Seq-Well RNA-seq Computational Pipeline and Analysis

Raw sequencing data was converted to demultiplexed FASTQ files using bcl2fastq2 based on Nextera N700 indices corresponding to individual arrays. Reads were then aligned to hg19 genome assembly and aligned using the Dropseq-tools pipeline on Terra (app.terra.bio). Data was normalized and scaled using Seurat R package v.3.1.0 (https://satijalab.org/seurat/), any cell with fewer than 750 UMIs or greater than 2,500 UMIs This cell-by-genes matrix was then used to create a Seurat object. Cells with any gene expressed in fewer than 5 cells were discarded from downstream analysis and any cell with at least 300 unique genes was retained. Cells with <20% of UMIs mapping to mitochondrial genes were then removed. These objects were than merged into one object for pre-processing and cell-type identification. The combined Seurat object was log-normalized to UMI+1 and applying a scale factor of 10,000. We examined highly variable genes across all cells, yielding 2,000 variable genes. Principal component analysis was applied to the cells to generate 100 principal components (PCs). Using the JackStraw function within Seurat, we identified significant PCs to be used for subsequent clustering and further dimensionality reduction. For 2D visualization and cell type clustering, we used a Uniform Manifold Approximation and Projection (UMAP) dimensionality reduction technique and with “min dist” set to 0.5 and “n neighbors” set to 30. To identify clusters of transcriptionally similar cells, we employed unsupervised clustering as described above using the FindClusters tool within the Seurat R package with default parameters and k.param set to 10 and resolution set to 0.5.nDifferential expression analysis between the negative and positive groups of the same CD8^+^ T-cells was performed suing the Seurat package FindAllMarkers in Seurat v3 (setting “test.use” to bimod). For each cluster, differentially expressed (DEGs) were generated relative to all of the other cells. Gene ontology and pathway analyses from DEGs were performed using Ingenuity Pathway Analaysis (IPA) which supports statistical analysis and visualization profiles for genes and gene clusters.

### SMART-Seq2 Whole Transcriptome Amplification and RNA Sequencing

Libraries were prepared using a modified SMART-Seq2 protocol as previously reported [83, 84]. In brief, based on FACS analysis (BD FACSAriaTM), single cells of Tetramer^+^ (HIV or CMV-specific), and tetramer- cells (bulk/total CD8^+^ T-cells), were sorted into wells onto of 96-well plates with lysis buffer, which contained 1 μl 10 mM dNTP mix, 1 μL 10 μM oligo dT primer, 1.9 μl 1% Triton X-100 (Sigma) plus 0.1 μl 40 U/μl RNase Inhibitor. The sealed plates were stored frozen at −80°C. Thawed plate is incubated for 3 min at 72°C and placed on ice. Next, SMART-Seq2 Whole Transcriptome Amplification (WTA) was performed: 7 μL of RT mix was added to each well and RT was carried out; then, 14 μL of PCR mix was added to each well and PCR was performed. Thereafter a cDNA clean-up was performed using 0.6x and 0.8x volumes of Agencourt AMPure XP SPRI beads (Beckman Coulter) to eliminate short fragments (less than 500 bp). Libraries was then quantified using a Qubit dsDNA HS Assay Kit (Life Technologies). Library size and quality were measured by Bioanalyzer using a High Sensitivity DNA Analysis Kit (Agilent Technologies). Sequencing libraries were prepared from WTA product using Nextera XT (Illumina). After library construction, a final AMPure XP SPRI clean-up (0.8 volumes) was conducted. Library concentration and size were measured with the KAPA Library Quantification kit (KAPA Biosystems) and a Tape Station (Agilent Technologies), respectively. Library were then constructed using Nextera XT DNA library preparation kit (Illumina FC-131-1096) using index primer. After library construction, sequences were purified using a 0.8X SPRI ratio yielding library sizes with an average distribution of 500-750 pb in length as determined using an Agilent hsD5000 Screen Tape System (Agilent Genomics). Finally, samples were sequenced on a NextSeq500 (30 bp paired-end reads) to an average depth of 5 million reads. Reads were aligned to hg19 (Gencode v21) using Top Hat [85] and estimated counts and transcripts per million (TPM) matrices generated using RSEM [86]. Any samples with fewer than 5×10E5 or more than 6×10E6 aligned reads or fewer than 10,000 uniquely ex-pressed genes were removed from subsequent analysis.

### TCR sequencing

A fraction of the WTA product was further amplified with V and C primers (both *α* and *β*) for TCR amplification. Followed by a second PCR reaction incorporating individual barcodes in each well. The samples are combined, purified and sequenced using Illumina Miseq with pair-end reads. For TCR sequences, the CDR3 nucleotide sequence are extracted and translated.

### SMART-Seq2 RNA-sequencing Data Analysis

Cells with a minimum of 2000 genes, maximum of twenty percent mitochondrial genes were included in downstream analysis run using Scanpy version 1.4.7 (scanpy.readthedocs.io) package [87]. PCA was run the log-normalized expression values of the top 1500 variable genes determined with the scanpy.preprocessing.highly variable genes function with inputs, (flavor=’seurat’, batch key=”patient”). Visualization using tSNE was run on top 10 principal components with perplexity 10 and learning rate 200. Preprocessing and clustering using Louvain clustering with resolution 0.7 were performed. Marker genes evaluated using rank genes groups function using method “t-overestem-var”. Top thirty genes were evaluated and ribosomal and mitochondrial genes were removed from gene lists. Plotting and visualization with matplotlib [88] and seaborn [89] libraries. Gene set scoring was run with the score genes function with parameter “use raw=True” using genes shared between reference gene lists and the transcriptional data. Differences in gene set score between groups was evaluated with a Kruskal Wallis test as implemented in scipy.stats.

The log of normalized MFI values for each channel associated with cells sequenced with SmartSeq2 were split into high and low bins using a Gaussian mixture model with two components using scikit-learn [90].

TCR alignment was executed using MiXCR [91]. Clones represented in two or more single cells were considered to be expanded for downstream analysis. The Scirpy package was used for further downstream analysis [92].

### Quantification and Statistical Analysis

Graphs were plotted using Prism 9.2.0 (GraphPad Inc.) Difference between groups were analysed using Mann Whitney U-test or ANOVA multiple comparisons test (two -sided), with a p value ¡0.05 considered to be statistically significant. If any other specific test used, it has been stated in the figure legends. The values of n refer to the number pf participants used in the study.

## Acknowledgements

We wish to thank the participants for their contribution to this study, the staff at the Africa Health Research Institute (AHRI) and associated medical and hospital clinical staff at Stanger and Addington Hospital in KZN. HNK is supported by the Wellcome Trust (202485/Z/16/Z). AL is supported by the Wellcome Trust (210662/Z/18/Z). This work was supported through the Sub-Saharan African Network for TB/HIV Research Excellence (SANTHE), a DELTAS Africa Initiative (grant DEL-15-006). The DELTAS Africa Initiative is an independent funding scheme of the African Academy of Sciences (AAS) Alliance for Accelerating Excellence in Science in Africa and supported by the New Partnership for Africa’s Development Planning and Coordinating Agency (NEPAD Agency) with funding from the Wellcome Trust (grant 107752/Z/15/Z) and the United Kingdom government. HNK and AS were supported by SANTHE. AKS was supported, in part, by the Searle Scholars Program, the Beckman Young Investigator Program, the NIH (5U24AI118672), a Sloan Fellowship in Chemistry, and the Bill and Melinda Gates Foundation. The views expressed in this publication are those of the authors and not necessarily those of AAS, NEPAD Agency, Wellcome Trust, or the United Kingdom government.

## Author contributions

RF performed experiments and prepared the manuscript. SN performed transcriptional analysis. OEA, SWJ, AS, AN, and JG contributed to experimental work. NM, DR and FK coordinated human sample collection. SB contributed with reagents. FA, JZP, ALS, RB, KM, WK contributed surgical human tissue samples. TN and PG contributed samples. SN, TdO, TN, PG provided intellectual input. AKS and BB supervised data analysis and provided intellectual input. HNK conceptualized andsupervised the work with intellectual input from SN, AKS and AL.

