## Supplementary Information for "HIV-specific CD8^+^ T-cells in tonsils express exhaustive T_RM_-like signatures"

Supplemental information includes 5 figures (Figure S1-S5) and 14 tables and can be found within this article. Supplemental tables can be found online at https://sid.erda.dk/sharelink/bStA6KGKwO

**Table S1** List of differentially expressed genes of $HIV^+$ $CD8^+$ T cells between Blood and Tonsil.

**Table S2A** The patient information and the responses from each patient sequenced.

**Table S3** List of differentially expressed genes between of HIV and CMV-specific $CD8^+$ T cells from scRNA-seq .

**Table S4** List of differentially expressed genes between $CD8^+CD69^+$ and $CD8^+CD69^+$

**Table S5** List of differentially expressed genes between $CD8^+CD103^+$ and $CD8^+CD103^+$

**Table S6** List of differentially expressed genes between $CD8^+CD127^+$ and $CD8^+CD127^+$

**Table S7** List of differentially expressed genes between $CD8^+$PD-$1^+$ and $CD8^+$PD-$1^+$

**Table S8** List of significant genes up-/downregulated between Viremic and Controller

**Table S9** Gene Set Analysis Results Using Ingenuity Pathway Analysis

**Table S10-S12** List of differentially expressed genes between clonotype expanded and unexpanded.

**Table S13** TCR$\alpha$ and TCR$\beta$ chain reconstruction from full-length single-cell transcriptome of every single cell.

**Table S14** Resource Table

### Supplementary Figures

Figure S1 related to Figure 1

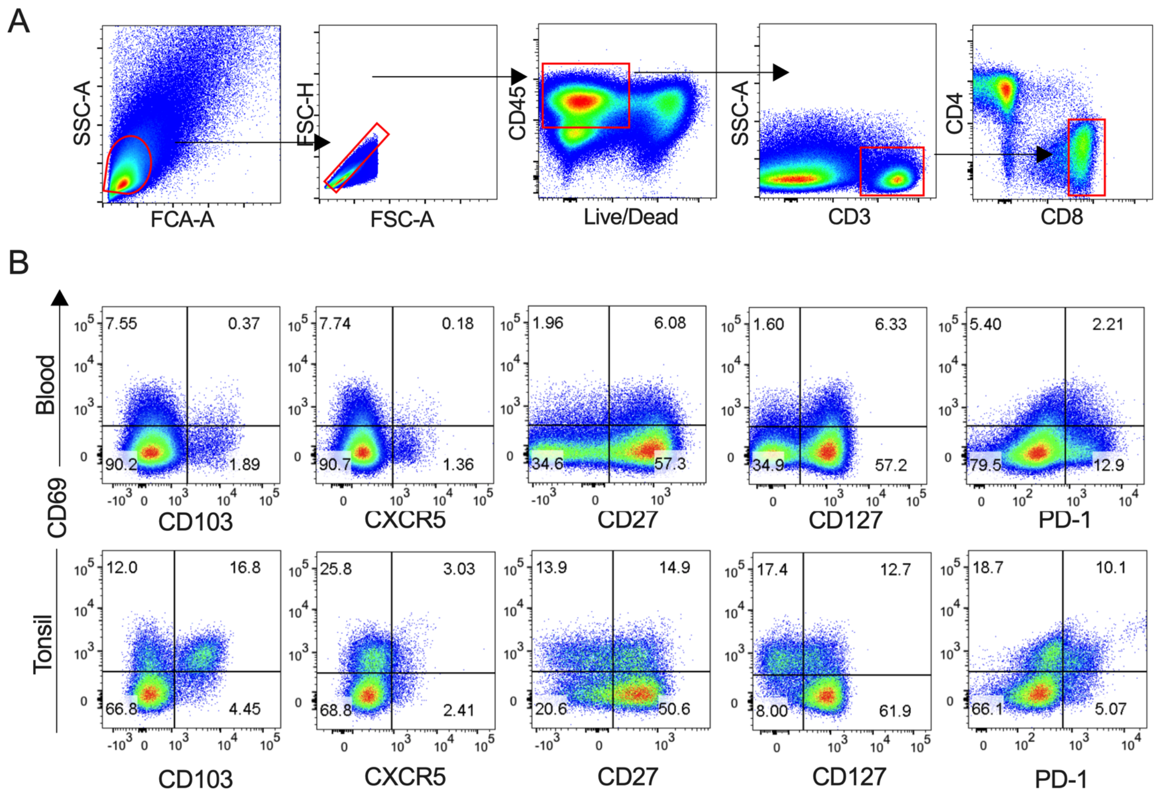


**Fig S** 1: **Supplementary figure 1. A** Representative flow cytometry plots (FACS) showing gating of tonsil CD8$^+$ T-cells. **B** Representative FACS plots of CD8$^+$ T cells as CD69+ expression versus CD103, CXCR5, CD27, CD127, and PD-1 in blood (top row) and tonsil (bottom row).

Figure S2 related to Figure 2

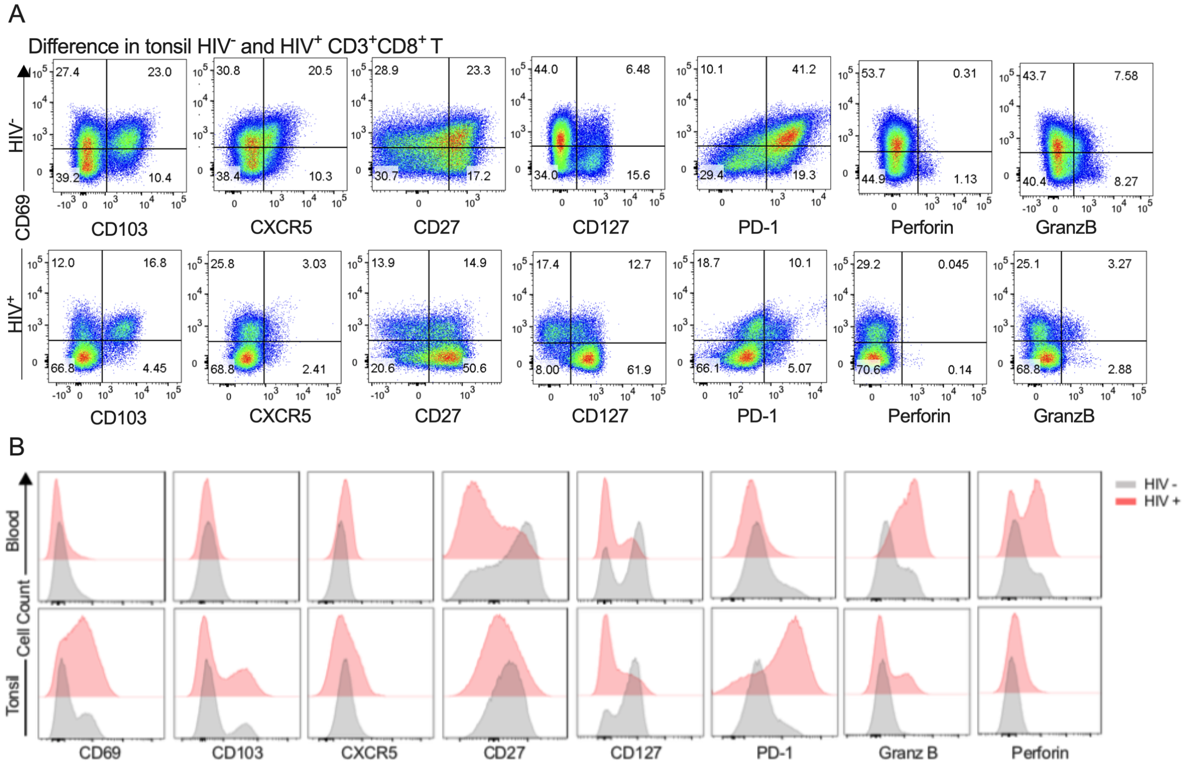


**Fig S** 2: **Supplementary figure 2.**. **A**. Representative flow cytometry plots (FACS) showing gating of tonsil CD8$^+$ T-cells from HIV- tonsil (top) and HIV+ tonsils (bottom) with FACS plots of CD8$^+$ T cells as CD69+ expression versus CD103, CXCR5, CD27, CD127, PD-1, perforin and granzyme B. **B**. Same as in A but showing histogram overlays for blood (top) and tonsil (bottom) CD8$^+$ T-cells.

### Figure S3 related to Figure 3

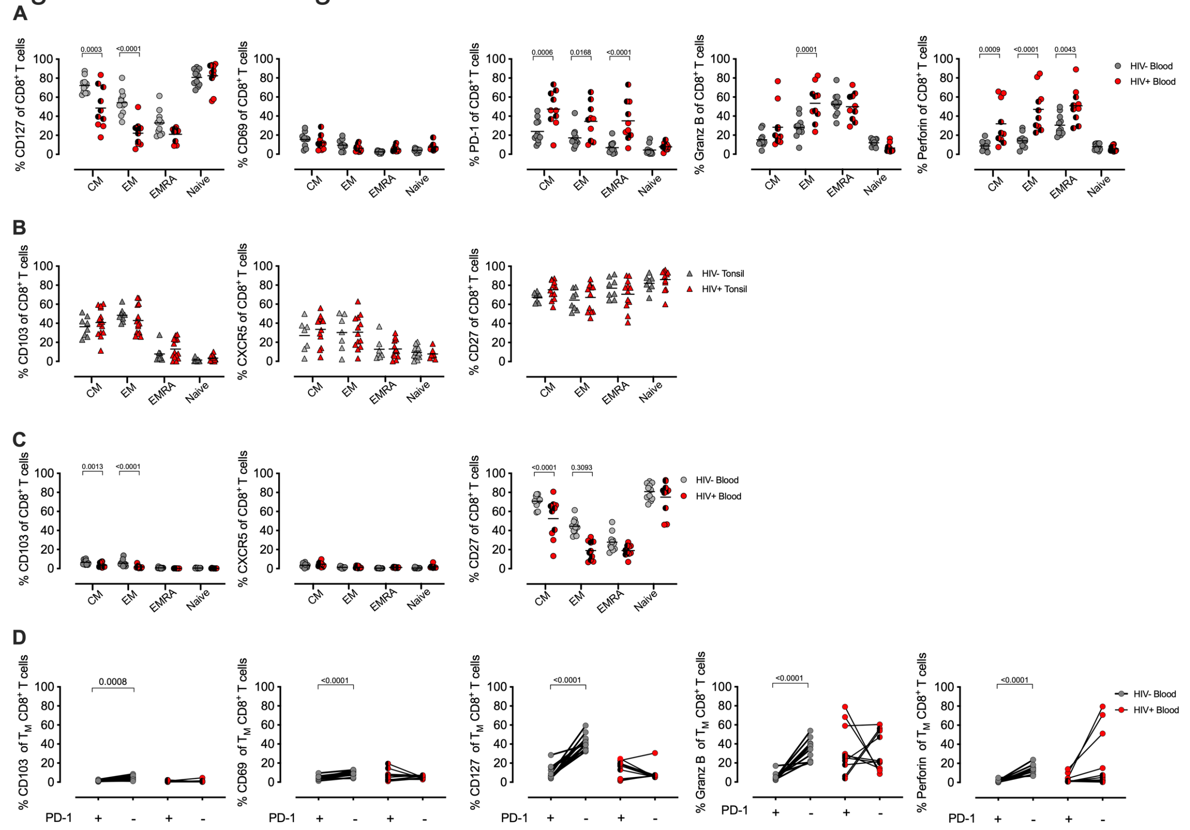


**Fig S 3: Supplementary Figure 3. CD8$^+$ T-cell memory profiling in blood and tonsils during HIV infection.**. **A**. Distribution of blood central memory (TCM), transitional memory (TEMRA), effector memory (TEM) and naïve subsets within CD127, CD69, PD-1, granzyme B and perforin expressing CD8$^+$ T-cells cumulative for all study participants in HIV- (grey) and HIV+ (red) with half black symbols HIV viremic participants. **B**. Same as in A but showing data from tonsil CD8$^+$ T-cells. **C**. Same as in A but showing data from blood CD8$^+$ T-cells for CD103, CXCR5 and CD27. D. P-values calculated using ordinary one-way ANOVA with horizontal bars representing median values with the level of significance indicated above. **D**. The frequency of CD103, CD69, CD127, perforin, and granzyme B (Granz B) cells measured on PD-1+ (left) and PD-1- (right) CD8$^+$ T-cells from blood in HIV+ (red) and HIV- (grey) individuals. P-values calculated using Paired Student's t test. Horizontal bars represent median values.

### Figure S4 related to Figure 5

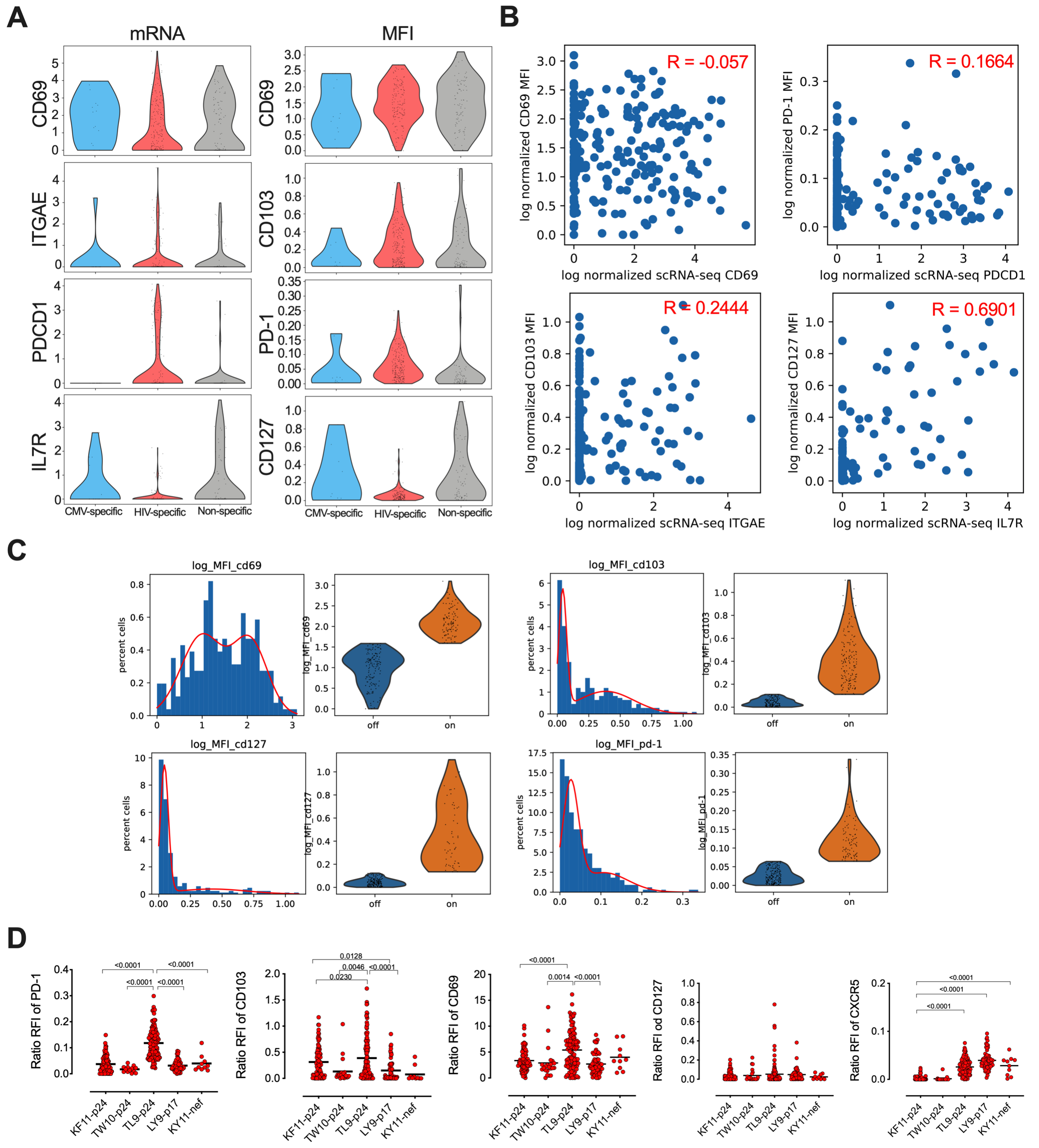


**Fig S** 4: **Supplementary figure 4. A**. Violin plots displaying mRNA expression from scRNA-Seq (left) and protein matched normalized MFI from flow cytometry (right) for matched single cells for each gene/protein with CMV, HIV and 'non-specific' CD8$^+$ T-cells indicated below. **B**. Correlation between mRNA expression from scRNA-Seq and normalized protein MFI from flow cytometry for matched single cells from A. **C**. MFI of histograms and the corresponding violin plots for single cells for CD69, CD103, CD127 and PD-1 expression in HIV-specific CD8$^+$ T-cells with on/off indicated and defined as either not expressed (0.0) or expressed (1.0) with cells ordered by high/low normalized MFI Genes ordered by hierarchical clustering. Select genes labelled in plot, full gene lists in Table S4-S7. **D**. Frequency of relative fluorescence intensity (RFI) of PD-1, CD103, CD69, CD127 and CXCR5 in HIV-tetramer specific CD8$^+$ T-cells.

Figure S5 related to Figure 6

**A**

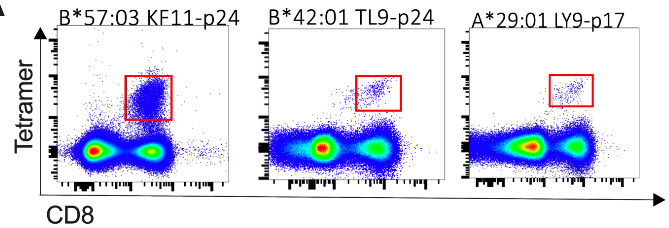


**B** Table of TCR alpha and beta clonotype sequence

27-14-0150 A* 42:01 TL9-p24

| TRAV<br>TRBV | CDR3α<br>CDR3β | Freq (%) |
|---|---|---|
| TRAV26-1<br>TRBV12-3 | CIVRGLNNAGNMLTF<br>CASSLGANPLHF | 33.3 |
| TRAV26-1<br>TRBV12-3 | CIVRGLNNAGNMLTF<br>CASSLGPNTIYF | 19.0 |
| TRAV13-1<br>TRBV12-3 | CAATNAGNNRKLIW<br>CASSFSKNTEAFF | 19.0 |
| TRAV26-1<br>TRBV12-3 | CIVRALNNAGNMLTF<br>CASSLGANTIYF | 9.5 |
| TRAV8-4<br>TRBV12-3 | CAVNSGGYQKVTF<br>CASSGTFYGYTF | 4.8 |
| TRAV26-1<br>TRBV12-3 | CIVRAPNNAGNMLTF<br>CASSLSINPVFF | 4.8 |
| TRAV26-1<br>TRBV12-3 | CIVRALYNAGNMLTF<br>CASSLGANTIYF | 4.8 |
| TRAV17<br>TRBV10-3 | CATDAPSGTYKYIF<br>CAAKDGGGTDEQFF | 4.8 |

27-14-0010 A* 29:01 LY9-P17

| TRAV<br>TRBV | CDR3α<br>CDR3β | Freq (%) |
|---|---|---|
| TRAV17<br>TRBV9 | CATGGFKTIF<br>CASSVGQYMNTEAFF | 54.2 |
| TRAV17<br>TRBV20-1 | CATGGFKTIF<br>CSATLNWGFF | 12.5 |
| TRAV19<br>TRBV20-1 | CALRSYSTLTF<br>CSATLNWGFF | 8.3 |
| TRAV17<br>TRBV20-1 | CATGGFKTIF<br>CSVTDYGYTF | 8.3 |
| TRAV9-2<br>TRBV6-2 | CALRGGATNKLIF<br>CASLRDYEQYF | 4.2 |
| TRAV20<br>TRBV19 | CAVGNNAGNMLTF<br>CASSIFGELFF | 4.2 |
| TRAV1<br>TRBV9 | CALRSYSTLTF<br>CASSVGQYMNTEAFF | 4.2 |
| TRAV19<br>TRBV19 | CALRSYSTLTF<br>CASSIFGEQYF | 4.2 |

27-14-0149 B* 57:03 KF11-P24

| TRAV<br>TRBV | CDR3α<br>CDR3β | Freq (%) |
|---|---|---|
| TRAV14<br>TRBV7-6 | CAMRESISSGSARQLTF<br>CASSHLGGGTDTQYF | 74.4 |
| TRAV14<br>TRBV7-6 | CAMRESISSGSARQLTF<br>CASSPPGTQYF | 20.9 |
| TRAV14<br>TRBV6-6 | CAMRESVSSAAGNKLTF<br>CAS RTGTPPYEQYF | 2.3 |
| TRAV14<br>TRBV20-1 | CAMRESISSGSARQLTF<br>CSATLNWGFF | 2.3 |

**C**

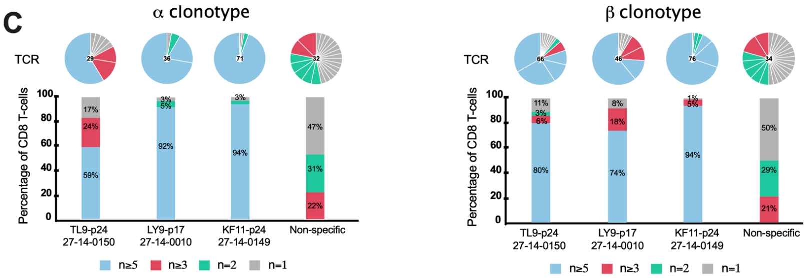


**D**

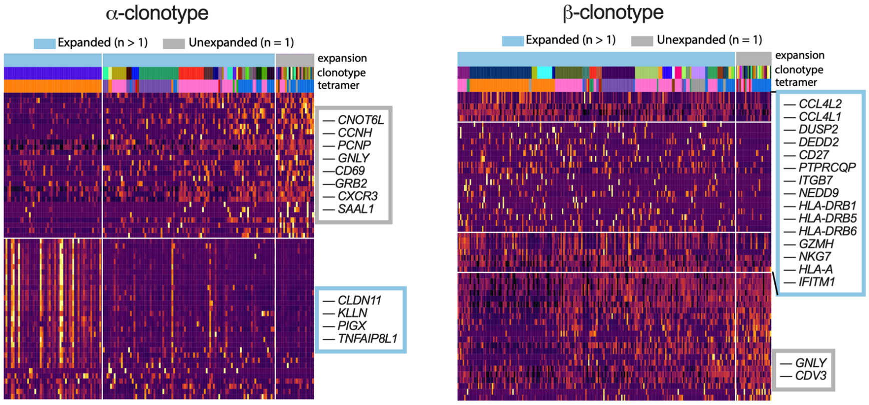


**Fig S 5**: **Supplementary figure 5.**. **A**. Flow plots showing KF11-p24 (B*57:03), TL9-p24 (B*42:01) and LY9-p17 (A*29:01) tetramer specific CD8$^+$ T-cells in tonsils. **B**. TCR and clonotype frequency. **C**. The TCR and TCR chain distribution of KF11-p24, TL9-p24, LY9-p17 from 27-14-0149, 27-14-0150 and 27-14-0010, respectively or non-specific (Tet-) (see Table S2) with unique (n=1), duplicated (n=2), triplet (n 3) and clonal (n5) with bars colored in grey, green, pink and blue representing the fraction of cells belonging to groups of clonotypes with either 1, 2, 3-4 or ¿5 clonotypes, respectively. Pie charts above each bar illustrate the composition of every individual TCR D. Heatmap of z-scored gene expression of top differentially expressed genes (t-test) between Louvain clusters from scRNA-seq data of clonotype. **D**, Same as in C but for clonotypes. Cells grouped by expansion of clonotype, genes grouped by hierarchical clustering. Full gene lists in Table S11 and S12.
